# Quantifying HLA transcripts by genotype in chimeric mixtures at single-cell resolution

**DOI:** 10.1101/2025.09.12.675951

**Authors:** Sami B. Kanaan, Jason G. Underwood, Rula Green Gladden, Everett Fan, Shruti S. Bhise, Monica S. Thakar, Carla A. Jaeger-Ruckstuhl, Jeffrey Stevens, Ashley N. Gray, Stanley R. Riddell, Marie Bleakley, Soheil Meshinchi, Scott N. Furlan

**Affiliations:** Translational Science and Therapeutics Division, Fred Hutchinson Cancer Center, Seattle, WA, USA; Pacific Biosciences of California, Menlo Park, CA, USA; Department of Pediatrics, University of Washington, Seattle, WA, USA; Clinical Research Division, Fred Hutchinson Cancer Center, Seattle, WA, USA; Seattle Children’s Research Institute, Seattle, WA, USA; Cancer and Blood Disease Institute, Pediatric Transplantation and Cellular Therapy, Children’s Hospital Los Angeles (CHLA), Los Angeles, CA, USA; Department of Medicine, University of Washington, Seattle, WA, USA; Department of Genome Sciences, University of Washington, Seattle, WA, USA; Brotman Baty Institute for Precision Medicine, Seattle, WA, USA

## Abstract

Gene products from the highly variable major histocompatibility locus, including HLA, are essential for self-recognition and immune surveillance of malignancy. Following allogeneic hematopoietic cell transplantation (alloHCT), genetic and epigenetic alterations in HLA can drive disease recurrence, making precise HLA assessment critical for determining future therapy. However, current methods lack the sensitivity to quantify HLA transcripts at the single-cell level, limiting their clinical utility. We introduce *scrHLA-typing*, a novel technique that accurately identifies and quantifies HLA transcripts in single cells using long-read sequencing. When applied to samples from patients with post-transplant relapse, scrHLA-typing successfully detected HLA allele-specific expression, across a range of levels of donor-recipient chimerism, at clinically actionable levels. By characterizing allele expression in residual leukemia cells, our assay identified differences in expression patterns among patients. This capability highlights scrHLA-typing’s potential to improve risk stratification and guide the selection of appropriate salvage therapies, enhancing personalized treatment strategies after relapse.

## MAIN

The human leukocyte antigen (HLA) genes reside in the extended major histocompatibility complex (MHC) region on chromosome 6p21.3, one of the most gene-dense and polymorphic stretches of the human genome^1^. By April 2025, about 42,584 HLA alleles had been catalogued across 46 clinically relevant genes in this region^2,3^. Variants in these loci account for more heritable disease than all other loci combined^4,5^. By presenting peptide antigens to T cells, HLA molecules are central to recognizing self vs. non-self, and thus to the adaptive immune response against pathogens, alloantigens, and neoantigens in cancer^6^.

In allogeneic hematopoietic cell transplantation (alloHCT), the effects of patient and donor HLA disparity on graft rejection, graft-vs.-host disease, and survival post-HCT are well established^7,8^. Importantly, when used as consolidation therapy for acute leukemias, alloHCT confers a beneficial graft-vs.-leukemia effect, in which HLA molecules play a crucial role^9^. Indeed, a frequent mechanism of immune escape and relapse post-alloHCT involves the loss of HLA surface expression on leukemic cells^10,11^. This alteration can result from epigenetic silencing of HLA or downregulation of regulators such as the CIITA^10,12^. Alternatively, a loss of HLA heterozygosity (LOH) has been well documented in mismatched alloHCT in the direction of HLA mismatch^13,14^, and even observed in matched alloHCT^15^. Generally, LOH of HLA is a well-described phenomenon in cancer, including outside the alloHCT context^16^, highlighting a broad link between allele-specific HLA loss and cancer progression.

Current genomic methods for HLA typing are predominantly performed on DNA obtained from bulk cell populations. While computational methods exist to infer HLA types and expression levels from bulk^17,18^ and single-cell RNA sequencing data^19–21^, no method currently enables the quantification of HLA molecules in complex mixtures. This limitation is particularly evident in settings such as post-alloHCT, where three or more alleles per gene can coexist at varying proportions—and often disproportionate levels. The high degree of homology among HLA genes and alleles (some differing by as little as a single nucleotide) introduces significant challenges, including variant call errors and alignment bias. These issues render traditional short-read sequencing pipelines inappropriate^22^ for use after transplantation. Consequently, most sequencing-based HLA genotyping approaches rely on statistical imputation of single nucleotide polymorphisms in linkage disequilibrium with known alleles^23^, or by alignment to a reference of all known alleles, subsequently narrowing down to one (homozygous) or two (heterozygous) alleles per gene by statistical inference^24–26^ or iterative realignment to a personalized reference^27–29^, disregarding supernumerary alleles as biologically or methodologically implausible.

We hypothesized that long-read sequencing of HLA transcripts in single cells could resolve the HLA landscape in individual cells, especially within complex mixtures such as those encountered after alloHCT. To this end, we developed <u>s</u>ingle-<u>c</u>ell <u>R</u>NA-based <u>HLA</u> <u>typing</u> (scrHLA-typing): a novel method that combines molecular enrichment of single-cell barcoded HLA full transcripts with long-read sequencing to enhance allele typing accuracy. A dedicated computational pipeline performs genotype prediction, error-filtering, and single-cell quantification accommodating complex allelic multiplicity. Applying scrHLA-typing to bone marrow aspirates (BMAs) from patients with relapsed leukemia following alloHCT, we demonstrate that this approach robustly quantifies differential HLA allele expression across diverse cell populations, providing clinically actionable insights.

## RESULTS

### Designing the scrHLA-typing workflow

Our technique relies on the generation of a single-cell barcoded whole transcriptome amplified pool (the cDNA pool) from a sample of cells, in which each cDNA molecule includes a cell-barcode (CB) sequence (to mark single-cell provenance) and a unique molecular identifier (UMI) sequence (to quantify and eliminate PCR duplication events). We crafted a panel of biotinylated DNA oligonucleotides (hybridization probes) that specifically recognize common regions of coding DNA sequence of classical and non-classical (including pseudogenes) HLA alleles in the IMGT/HLA database^3^ (Fig. 1a). Although optimized for 10X Genomics-derived cDNA, any single-cell barcoded whole transcriptome should be compatible. After hybridization capture, HLA fragments were then re-amplified and prepared specifically for PacBio long-read sequencing (Fig. 1a).

**Fig. 1.**
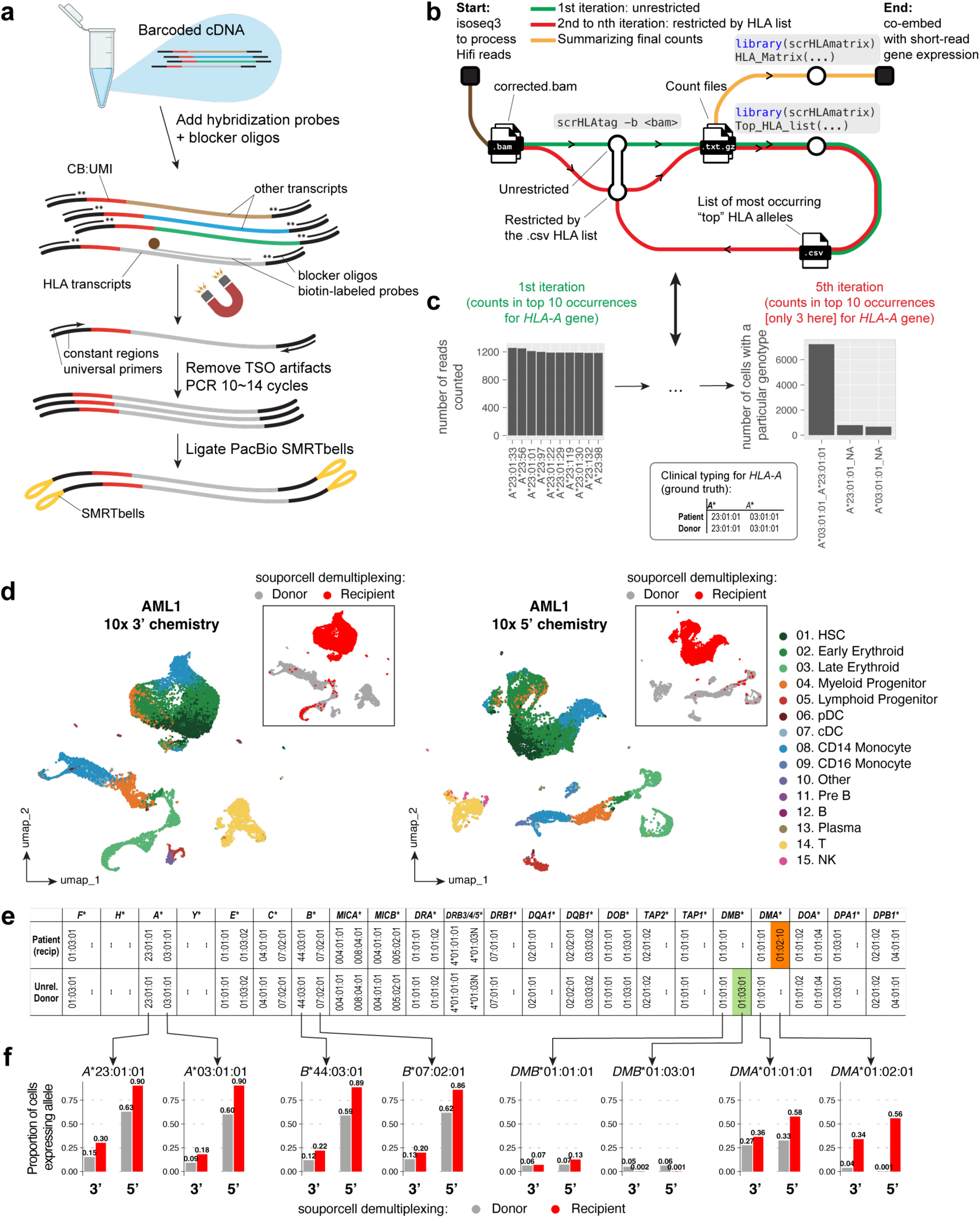
Method schematization and comparison across single-cell capture chemistries. **a**, Single-cell barcoded cDNA pools are combined with custom-designed biotinylated hybridization probes and blocking oligos. Enriched HLA transcripts are depleted of template switch oligo (TSO) priming artifacts, PCR-amplified, and prepared for long-read sequencing. **b**, PacBio HiFi reads are processed with the IsoSeq pipeline to generate deduplicated BAM files, serving as the input for the first ‘unrestricted’ scrHLAtag iteration (green line). scrHLAmatrix predicts a list of top HLA candidates, which is then provided back to scrHLAtag for one or more ‘restricted’ iterations (red line) until a final candidate list is obtained. Count files from the final round are summarized into a sparse numeric matrix. **c**, Left: Top 10 most numerous alleles in the example of *HLA-A* generated in the first unrestricted scrHLAtag iteration for a sample from patient ‘AML1’; right: Top 10 most numerous genotype configurations in the example of *HLA-A* generated in the final unrestricted scrHLAtag iteration for the same sample from patient ‘AML1’. **d**, UMAP embedding of profiles colored by cell type and genetic demultiplexing result (inset) from patient ‘AML1’ captured using 3-prime and 5-prime chemistries. **e**, Genotyping result of donor and recipient cells from patient ‘AML1’ using scrHLA-typing. **f**, Fraction of cells expressing each allele, stratified by donor vs. recipient classification, illustrated for two genes without donor– recipient mismatches (*HLA-A*, *HLA-B*) and two genes with mismatches (*HLA-DMA*, *HLA-DMB*).

For data analysis, we developed a pipeline with two components. ‘scrHLAtag’, a command line tool, aligns reads to either the full or partial list of the IMGT/HLA reference (i.e. when HLA types are known). When HLA types are not known *a priori*, a ‘best-guess’ list is iteratively generated combining scrHLAtag with ‘scrHLAmatrix’, an R package that detects HLA patterns, summarizes calls per cell, filters errors and produces data objects that integrate into current short-read analysis pipelines (Fig. 1b). In patient samples used throughout this study, scrHLAmatrix-to-scrHLAtag typically converged on the putative HLA genotypes after 5 to 9 iterations (Fig. 1c,e; Suppl. Tables S1, S2).

For our first experiment, we performed scrHLA-typing on a gradient centrifuged BMA sample from an 11-year-old male patient with CD34+ acute myeloid leukemia (AML), collected 783 days after HLA-matched unrelated donor myeloablative alloHCT when low level relapse had been confirmed clinically using multiparameter flow cytometry. To boost numbers of leukemic blasts, we also captured a fraction of the sample after performing CD34+ immunomagnetic enrichment. We then performed single-cell RNA droplet capture of both unenriched and CD34+ cells using both 10x Genomics 5’ and 3’ single-cell chemistries, followed by whole transcriptome short-read sequencing.

To establish ‘ground truth’ we used souporcell, a genetic demultiplexing algorithm capable of distinguishing donor vs. recipient cells using variants present in the aligned reads^30^. We then used viewmastR^31^ to annotate cell types according to a known reference, in our case, a well annotated atlas of healthy bone marrow mononuclear cells^32^. Putative leukemic ‘recipient’ cells annotated as hematopoietic stem cells, monocytes, and erythroid and myeloid progenitors in an overlapping continuum typical of dysregulated differentiation. In contrast, putative ‘donor’ cells followed an orderly hierarchical pattern of normal hematopoietic differentiation (Fig. 1d). To unequivocally confirm the identity of these cells using an approach previously described^33^, we designed biotinylated hybridization probes specific to two of the genes known to be mutated in the leukemia of this patient: *TP53* and *IDH1* (Suppl. Table S2). As expected, mutated *TP53* and *IDH1* were present only in leukemic recipient cells (Extended Data Fig. 1).

To generate HLA-to-CB count matrices that could be integrated with short-read profiles, we first filtered low quality reads (based on aligner scores; Extended Data Fig. 2), then removed PCR duplicates (with additional post-deduplication corrections; see Online Methods). Even after filtering for reads with low quality, we noted the presence of reads (with identical CB and UMI) that contained amalgam sequences made up of 2 HLA alleles. We attributed this phenomenon to a template-switching PCR artifact which has been described in sequencing applications^34^, but we hypothesize is more common when input libraries are less complex (Extended Data Fig. 3) such as when amplifying HLA sequences, where extensive polymorphism and inter-allele/inter-gene homology are defining features. As such, we adopted a deduplication strategy using ‘majority-wins’ approach. To evaluate the robustness of our computational strategy for HLA-based cell classification, we conducted *in silico* mixing experiments using combinations of 3 to 6 post-alloHCT patient samples (see Online Methods) and achieved consistently high classification accuracy (Extended Data Fig. 4).

We next asked how per-cell read coverage and erroneous allele calls (i.e., donor-specific alleles in recipient cells and vice-versa, recipient-specific alleles in donor cells) compared between the 2 capture chemistries. We found the proportion of cells expressing any HLA allele was higher in the 5’ captures, with intra-class molecular swapping events ∼45 times more common in 3’ captures (Extended Data Fig. 3d). This may be explained by the position of hypervariable HLA regions on the 5’ side (Extended Data Fig. 5). Finally, among the only 2 non-shared alleles in an otherwise perfectly HLA-matched donor-to-recipient alloHCT (Fig. 1e; Extended Data Fig. 6), the false positive rates (i.e., erroneous allele calls) were from 2 to 40-fold higher in 3’ vs. 5’ (Fig. 1f).

To broaden the potential usage of scrHLA-typing onto single nuclei preparations, we experimented on cDNA from a sample of human T cells that others had captured in a co-assay of single-cell RNA and ATAC^35^, in which cells were lysed/permeabilized to allow nucleus tagmentation and barcoding while simultaneously barcoding RNA at the 3’ end. These nuclei captures had a dearth of HLA transcripts and very few reads passed quality control (Extended Data Fig. 7). Given the superiority of pairing scrHLA-typing with 5’ RNA capture chemistry we performed subsequent experiments using 5’ barcoded samples with intact cells.

### Quantifying HLA in samples with disproportionate or more balanced chimerism levels

To assess scrHLA-typing across a range of disease burdens, we obtained samples from a 6-year-old patient with AML (AML4), collected post-haploidentical (donor source: father) myeloablative alloHCT at three timepoints. At day 90, when AML relapse was suspected clinically (0.04% burden by multiparameter flow cytometry), and at days 140 and 163, after overt clinical relapse (11.5% and 43%, respectively). Genetic demultiplexing results reflected clinical burden of disease (Fig. 2a). Remarkably, low level chimerism had no influence on the number of iterations needed for convergence (Extended Data Fig. 8 and Suppl. Table S1). Furthermore, scrHLA-typing results had high fidelity to genetic demultiplexing (Fig. 2).

**Fig. 2.**
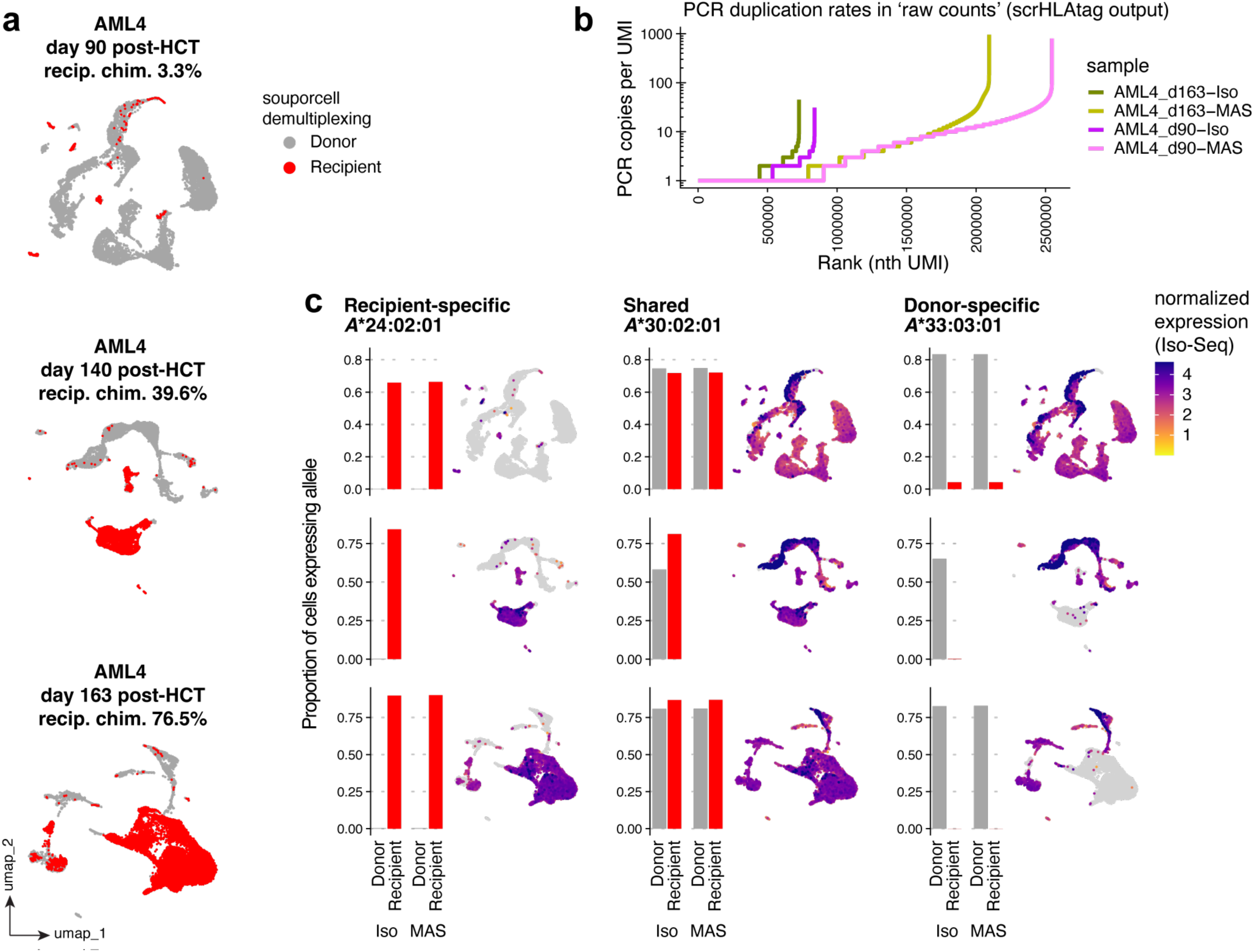
Quantifying HLA alleles longitudinally after post-transplant relapse. **a**, UMAP embedding colored by genetic demultiplexing result from patient AML4 BMMCs at days 90, 140, and 163 post-allo-HCT. **b**, PCR duplication rates in raw count data from scrHLAtag for 2 time points: ‘AML4 day 90’ and ‘AML4 day 163’, using either the standard IsoSeq PacBio method or the augmented MAS-seq method. **c**, Fraction of cells expressing each of the three HLA-A alleles, stratified by donor vs. recipient classification (souporcell), for each of the three ‘AML4’ time points. Normalized expression values for each allele are shown alongside as a quantitative measure.

Next, to establish whether increasing sequencing depth improved scrHLA-typing sensitivity, we employed ‘multiplexed arrays isoform sequencing’ (MAS-seq)^36^, in parallel with our standard ‘Iso-Seq’, on the samples ‘AML4 day 90’ and ‘AML4 day 163’ (Extended Data Fig. 9). Although MAS-seq increased raw read output 12–16 fold vs. Iso-Seq, PCR-duplication rates were considerably higher (Fig. 2b). Cell barcode coverage remained comparable with Iso-Seq (Extended Data Fig. 9c), normalized HLA expression patterns were similar (Extended Data Fig. 9e), and the ability to resolve alleles for each allogeneic entity and the resulting erroneous allele calls (false positives) were almost indistinguishable from Iso-Seq (Fig. 2c), with a global average false positive rate at 0.25% for 5’ chemistry captures (Extended Data Fig. 10). In all, results did not show significant benefit for using the higher throughput of MAS-seq across a range of donor chimerism levels, though it would still be considered for highly multiplexed future experiments.

### HLA transcript count correspondence with cell surface molecular expression

In an effort to assess whether quantification of HLA transcript correlated with protein level expression, (acknowledging the difficulty of resolving HLA at the protein level with current antibody technology), we evaluated pan *HLA-A*, *-DRA*, and *-E* on the cell surfaces using CITE-Seq^37^ for ‘AML4’ samples. To account for the well-described effect of cell state (steady vs. in-transition, long-term commitment vs. short-term adaptation, differentiation status, etc.) on mRNA-to-protein correlation^38–40^, we considered cell types as a proxy, and found heterogeneity in HLA allelic transcript-to-protein correspondence by cell type (Fig. 3a,b). In general, cell types clustered into 2 groups: exhibiting positive correlation (example in Fig. 3c) or exhibiting no significant correlation and sometimes negative correlation suggesting transcript-to-protein ‘signal delay’ or prolonged ‘persistence’ of stable protein without its encoding transcript (Fig. 3b). Notably, in some cell types, for example the recipient-specific late erythroid progenitors for the 3 timepoints (example in Fig 3.c) and the recipient-specific CD14+ monocyte-like cells for ‘AML4 day 90’, transcripts of one allele (e.g., the ‘shared’ allele) had slightly stronger correlation with surface molecule expression vs. the second allele, suggesting a stronger contribution of the former rather than the latter to protein formation. This analysis reveals that the correlation between HLA allelic transcript counts, and corresponding cell surface protein quantities is cell type-dependent, and occasionally allele-dependent. This underscores the complexity of mRNA-to-protein stoichiometry, influenced by cell state and lineage.

**Fig. 3.**
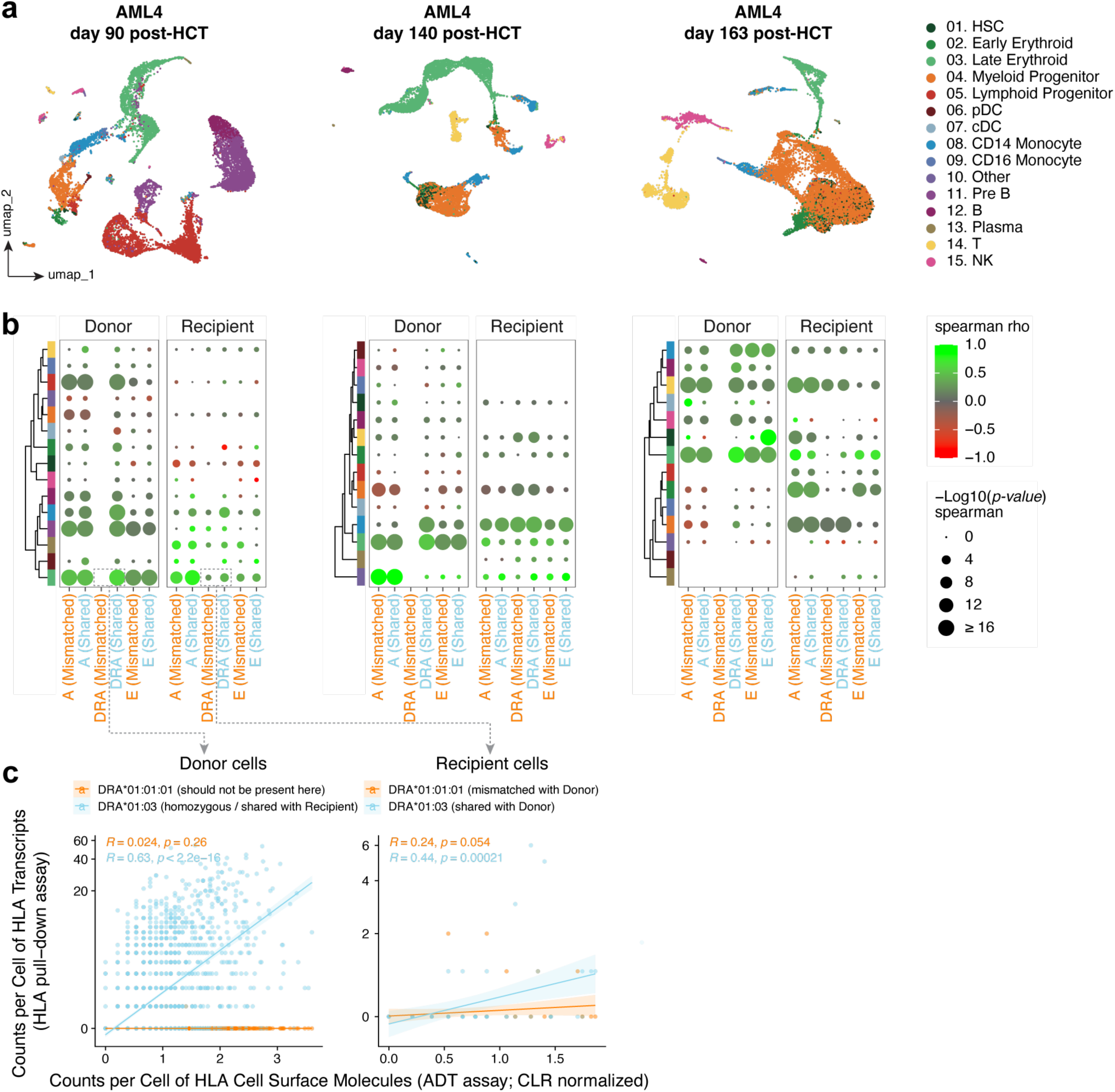
Correlation of HLA allele-specific transcript counts with cell surface HLA molecular expression. **a**, UMAP embedding of bone marrow mononuclear cell profiles of ‘AML4’ colored by cell type. **b**, The effect size (spearman rho) and statistical significance (*p*-values, spearman) across cell types for the relationship between per-cell counts of donor-specific (mismatched relative to the recipient) and shared HLA alleles in donor cells, and vice versa, recipient-specific and shared HLA alleles in recipient cells, with the corresponding cell-surface HLA molecules. **c**, Example correlations between transcript counts for *DRA*\*01:01:01 *and DRA*\*01:03 and cell-surface DRA molecule counts in the late erythroid subset from both recipient and donor cells of ‘AML4’ at day 90 post– alloHCT.

### Differential HLA allele expression is present in leukemia cells of some but not all patients post-alloHCT

Next, we asked whether mismatched HLA alleles (vs. shared) were differentially expressed longitudinally in ‘AML4’ (Fig. 4a,b). Given that persistent recipient T cell chimerism was present in ‘AML4’, we further subset the recipient cells into ‘malignant’ vs. ‘healthy’. Like in patient ‘AML1’, putative leukemia cells 1) exhibited evidence of disorganized differentiation (Extended Data Fig. 11a), 2) expanded in numbers over time (Extended Data Fig. 11b), 3) expressed genes implicated in cellular immortality^41^ (Extended Data Fig. 11c), and 4) expressed the AML-defining *KMT2A*::*MLLT3* fusion (Extended Data Fig. 12) clinically detected in this patient (cryptic fusion difficult to detect, negative by karyotype and fluorescence in situ hybridization cytogenetic methods, eventually characterized by the FusionPlex^42^ and UW-OncoPlex^43,44^ molecular assays).

**Fig. 4.**
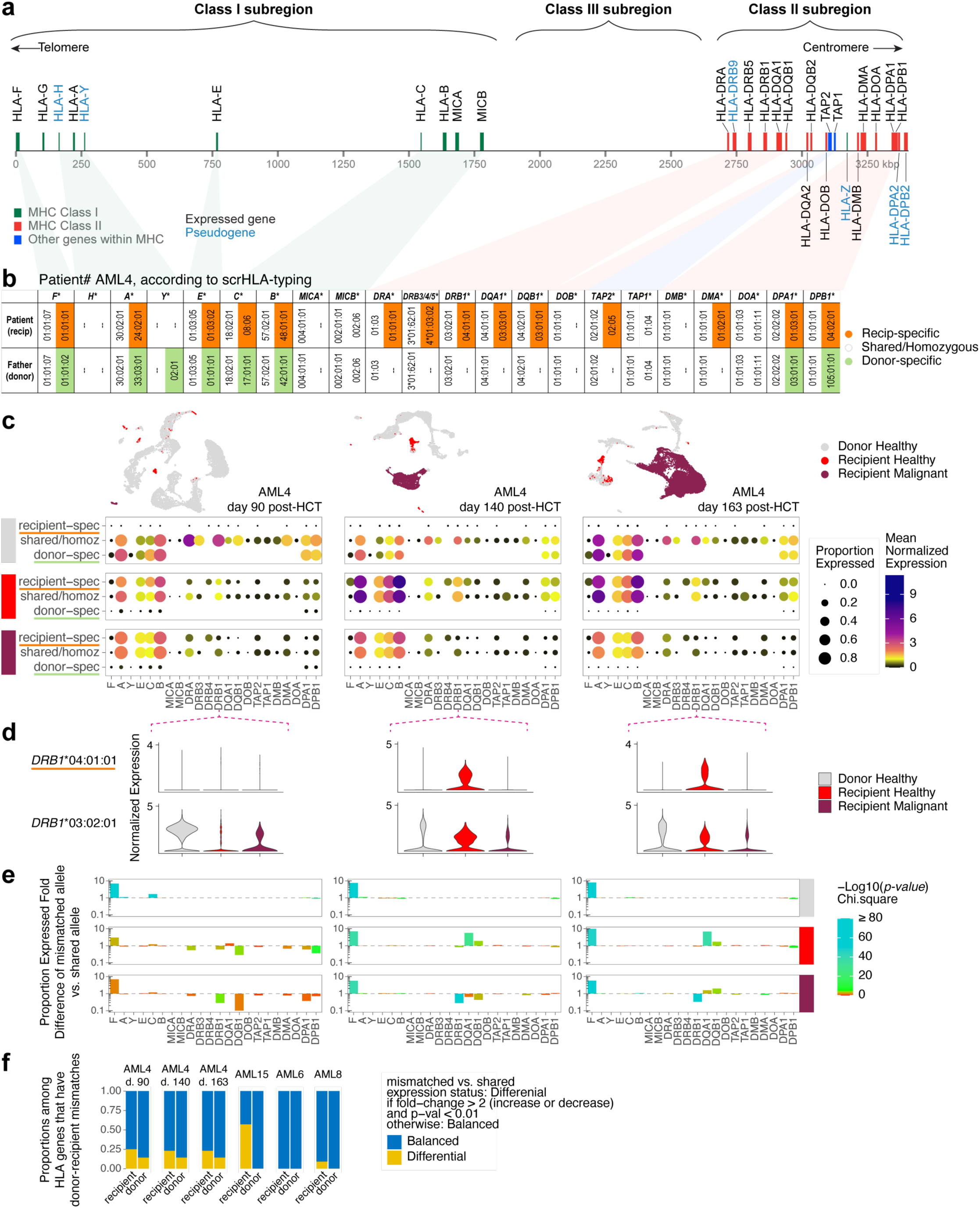
Differential Allele-Specific Expression (ASE) across the HLA haplotype in different groups of cells. **a**, HLA gene positions on chromosome 6 with coordinate ‘0’ corresponding to position 29,722,775 bp of the hg38 primary assembly (position of pseudogene *HLA-Y* was recently described in a ∼60,000 bp indel polymorphism [ref. Alexandrov, N., et al., HLA 102, 599-606 (2023)] between *HLA-A* and *HLA-E*). **b**, Genotyping of ‘AML4’ using scrHLA-typing. **c**, Mean normalized expression in the example of *HLA-DRB1*, for the recipient-specific (*DRB1*\*04:01:01) and the shared (*DRB1*\*03:02:01) alleles across the cell categories and timepoints. **d,** Example using *HLA-DRB1*, showing mean normalized expression of the recipient-specific allele (*DRB1*\*04:01:01) and the shared allele (*DRB1*\*03:02:01) across cell categories and time points. **e**, Fold change in the proportion of cells expressing the mismatched allele (under selective pressure) vs. the shared allele, along with statistical significance (Chi-square test), shown across cell categories and time points. **f**, Proportion of HLA genes exhibiting ‘balanced’ vs. ‘differential’ allele expression among genes with recipient–donor mismatches. ‘Differential’ expression is defined here as a fold change in the proportion expressed of mismatched vs. shared alleles > 2 (either increase or decrease) and a Chi-square p-value < 0.01.

In all 3 groups: donor healthy, recipient healthy (mostly lymphocytes), and recipient malignant, class II genes exhibited a greater degree of unbalanced allele-specific expression (ASE) than class I (Fig. 4c,e). Among leukemic cells, some recipient-specific class II alleles were downregulated vs. their shared counterpart. In particular, the recipient-specific *DRB1* allele (*DRB1*\*04:01:01) was significantly downregulated compared to the shared (*DRB1*\*03:02:01) in recipient malignant cells but not in recipient healthy cells (more obvious in later timepoints), suggesting a cell type-dependent rather than an allele intrinsic-dependent mechanism (Fig. 4d). Highly statistically significant unbalanced ASE in *DRB1* was evident early and remained consistent across the 3 timepoints post-alloHCT (Fig. 4e).

We tested 3 additional post-alloHCT AML patients for mismatched allelic downregulation within leukemic cells (Extended Data Fig. 13). Taking a similar approach to determine malignant cells as described above (using fusions if available; Extended Data Fig. 12), one patient, ‘AML15’, with recurrent disease post-haploidentical transplant (source: mother) had evidence of an erythroblastic leukemia (Extended Data Fig. 13a,b) that also exhibited unbalanced ASE (Extended Data Fig. 13c), strongest among class II alleles (Extended Data Fig. 13d). In other patients, unbalanced ASE was not appreciable in leukemic cells; patients ‘AML6’ and ‘AML8’ with relapse post-alloHCT (source: matched unrelated and single cord blood, respectively), had little differential allele expression per HLA gene across the haplotypes. For ‘AML8’, only the recipient-specific intracellular-expressed *HLA-DOA* was significantly downregulated vs. shared (Extended Data Fig. 13d).

Moreover, unlike recipient cells, the donor cells of all 4 patients generally maintained ‘balanced’ HLA expression across the haplotypes, in line with the understanding that cells from the donor maintain alloimmunologic pressure after alloHCT^45^. Furthermore, ASE of HLA in settings of donor-recipient mismatch was always more prevalent in recipient cells compared with donor cells (Fig. 4f) using thresholds of an absolute fold-change of 2, and p-value < 0.01. Altogether, our assay robustly detected ASE present in HLA genes. Some patients exhibited such differential expression in their leukemia cells, but others did not. We anticipate the ability to make this distinction among patients to become useful for the future of clinical decision-making after transplantation.

### Assessing mismatch patterns in HLA sequencing reads across single cells

We next asked whether scrHLA-typing could identify sequence variation in the coding region of HLA. Reads that mapped correctly to their corresponding IMGT/HLA reference sequence showed a high-fidelity rate, averaging a 99.7% match across all nucleotide positions, loci, and patient samples. Minor mismatches observed were evenly distributed across both loci and patient samples, likely attributable to technical artifacts from reverse transcription, PCR, and sequencing. An illustrative example of this low-level variation at the *HLA-A* locus in the ‘AML4 day 90’ sample is shown in Fig. 5a.

**Fig. 5.**
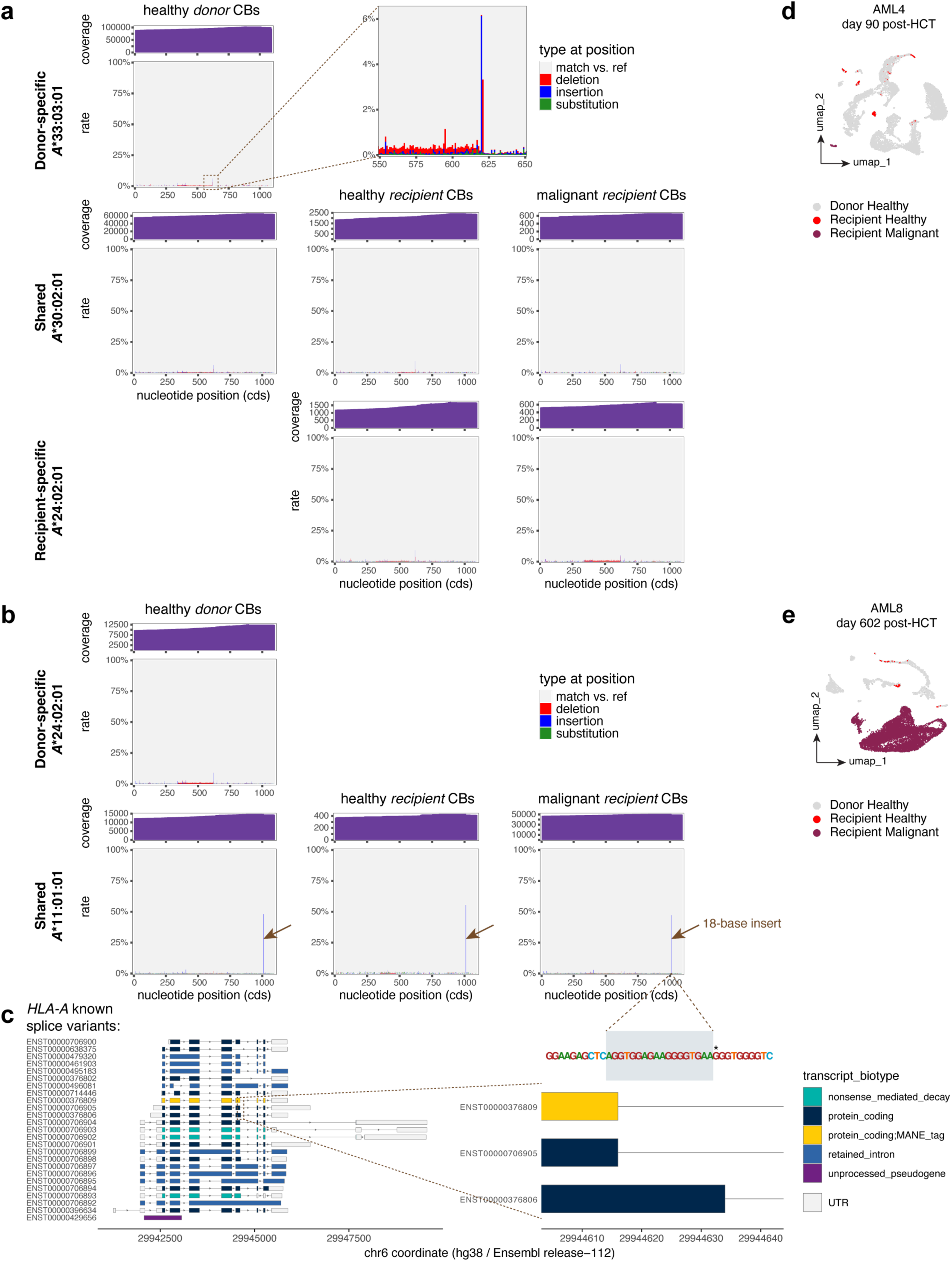
Mismatch patterns in HLA sequencing reads across single cells. **a,** Per-base mismatch rates in aligned reads, calculated relative to the assigned IMGT/HLA reference coding sequence (CDS), for *HLA-A* in ‘AML4’ at day 90 post–allo-HCT. Cells are classified by barcode (CB) as healthy donor, healthy recipient, or malignant recipient. Mismatch profiles are shown alongside sequencing coverage tracks (purple). For clarity, reads from erroneous allele calls (e.g., donor-specific alleles in recipient cells or vice versa, recipient-specific alleles in donor cells)—which typically display higher variant rates and markedly lower apparent coverage—are omitted. **b,** Per-base mismatch rates and coverage for *HLA-A* in ‘AML8’, highlighting a 3′-end insertion in *A\**11:01:01 affecting about half the reads (brown arrow), observed in healthy donor, healthy recipient, and malignant recipient cells. **c**, This 18-base insertion corresponds to a partially retained intron in some known *HLA-A* splice variants. The nucleotide at hg38 position 29,944,633 (marked with an asterisk) is a ‘G’ in the hg38 reference but an ‘A’ in the *HLA-A\**11:01:01 allele family. **d, e,** UMAP embeddings for ‘AML4 day 90’ (d) and ‘AML8’ (e), serving as cell-type keys for panels (**a**) and (**b**), respectively.

In contrast, a variant was identified in patient ‘AML8’, who was *HLA-A*\*11:01:01 homozygous and had received a transplant from an *A*\*11:01:01/*A*\*24:02:01 donor. Specifically, we observed an 18-base insertion at the 3’ end of the *A*\*11:01:01 reads, with a variant allele frequency of 47.4%. Our single-cell approach uniquely enabled us to evaluate this insertion across different cell populations; looking across healthy donor, healthy recipient, and malignant recipient cells, the insertion remained consistent at approximately that same frequency, unlikely to be disease-specific (Fig 5b). We further identified this insertion as a known alternative splice of *HLA-A* (Fig 5c), resulting in an in-frame addition of 6 residues (GGEGVK) at position 314 at the cytoplasmic domain start^46^, albeit with unknown consequence(s).

Collectively, these findings underscore the exceptional sensitivity of scrHLA-typing in detecting clinically relevant coding variations within HLA. Crucially, our method allows detailed analysis of HLA variation within specific cellular contexts, providing valuable insights with significant clinical implications for patients experiencing leukemia relapse post-transplant.

## DISCUSSION

We presented scrHLA-typing, a single-cell, long-read sequencing approach that resolves the full allelic complexity of HLA transcripts with three-field resolution in highly chimeric samples. By combining targeted hybridization capture, PacBio HiFi sequencing and an iterative genotyping-quantification pipeline, scrHLA-typing accurately calls and counts alleles including when multiple donor/recipient haplotypes coexist at disproportionate levels—a context where current bulk and short-read methods fail.

In post-alloHCT pediatric AML samples, scrHLA-typing correctly predicted classical class I and II HLA genotypes concordant with clinically-available donor/recipient genotyping and orthogonal genetic demultiplexing. Our assay retained its accuracy across extreme levels of chimerism and was able to quantify statistically significant cell type-specific ASE which were prevalent in malignant cells of some, but not all patients. This ability to discriminate meaningful ASE at single-cell resolution enables a new dimension of post-alloHCT monitoring with clinical implications for salvage therapy planning. Finally, our assay enabled high-fidelity detection of genetic variation within HLA exonic regions, though in this small cohort, no variants were identified that could be attributable to patients’ relapse.

A major advantage of our assay is its application of high-fidelity long-read sequencing at the single-cell level to resolve highly complex HLA loci—an advance over previous single-cell HLA studies^19–21^, which, while conceptually elegant were confined to short-read sequencing and limited to 2-field HLA resolution. To our knowledge, this is the first study to achieve in RNA three-field HLA genotyping and quantify alleles in complex chimeric mixtures such as those encountered in the post-alloHCT setting. The high base accuracy of our approach facilitates detection of somatic mutations in HLA, which are known to recur in cancer^25^. Such mutations can result from errors during DNA repair or replication^47–50^, but to be detectable, should involve a relatively small number of nucleotides. Indeed, longer genomic alterations, such as those resulting from homologous recombination^51^—the leading mechanism behind copy-neutral LOH of HLA—may appear as increased expression of the duplicated allele, thus indistinguishable from ASE-inducing epigenetic alterations; a potential limitation of our assay that warrants further study.

Additional limitations include reduced accuracy of HLA pattern-based classification of chimeric entities when the number of mismatched alleles between them is low (Extended Data Fig. 14). In such cases, however, the need for separate classification diminishes, as most alleles are shared and can be genotyped and quantified collectively. Finally, while we observed correlation between HLA transcriptional counts and cell surface protein expression in some cell types, the relationships were absent in others. This is consistent with the known complexity of mRNA–protein stoichiometry, which can be influenced by cell type, differentiation status, mRNA decay rates, protein synthesis and turnover kinetics, and temporal lags between transcription and translation, as well as prolonged protein persistence after new transcription or translation ceases— phenomena that differ between short-term adaptive states and more stable, long-term cellular processes^38–40,52,53^.

We anticipate that scrHLA-typing could have immediate clinical utility in risk-stratifying relapsed alloHCT patients prior to salvage therapy, by identifying likely mechanisms of immune escape and guiding therapeutic decisions. Critically, the assay can provide such insight even at low disease burden, when targeted interventions are most likely to succeed. Looking forward, assessing variability in HLA expression within leukemic cell clusters at diagnosis may also inform donor selection—prioritizing mismatches in alleles with consistently high and stable expression over those with lower or more variable expression—thereby optimizing graft-versus-leukemia potential from the outset.

## ONLINE METHODS

### Ethics statement

The study included pediatric patients who received alloHCT for acute leukemia, treated at the Seattle Children’s Hospital, and whose parents/guardians consented for collection of research study specimen and data collection SCH-B802. The Internal Review Board (IRB) of the Fred Hutchinson Cancer Center approved this study (IRB file #FHIRB0010627) which was conducted in accordance with institutional guidelines, the Declaration of Helsinki, and Title 45 United States Code of Federal Regulations, Part 46, Protection of Human Subjects.

### Specimen collection and processing

Bone marrow aspirates (BMA) were obtained from the pelvic bone of patients and were processed using density gradient centrifugation to isolate mononuclear cells. Any remaining red blood cells (RBCs) were depleted by incubating the mononuclear cells in an RBC-lysis buffer (NH_4_Cl 8.3 g/L; NaHCO_3_ 1 g/L; 0.04 g/L EDTA; filtered 0.22 um) for 10 min at room temperature, then washed in 1X PBS with 2% FBS. Cells could be viably cryopreserved in fetal bovine serum with 10% DMSO stored in liquid nitrogen for later thaw and use, or immediately used in the next processing steps. Cells were counted on a Nexcelom instrument to determine numbers and viability using ViaStain AOPI Staining (Nexcelom #CS2-0106-25mL). When viability < 80%, live cells were enriched using magnetic removal of dead cells (Miltenyi Biotec 130-090-101). When applicable, at most 10 million cells were enriched for cells with CD34 cell surface markers using either Miltenyi magnetic CD34 Microbead kit (Miltenyi Biotec #130-097-047) or EasySep human CD34 Positive Selection kit (Stemcell #17856) with an expected yield of ∼500,000 CD34-enriched cells. For patient ‘AML15’, instead of a CD34+ enrichment, cells with CD117 markers were enriched using Miltenyi magnetic CD117 Microbead kit (Miltenyi Biotec #130-091-332) in a decision based on the patient’s leukemia characteristics.

### Feature barcode cell staining

To profile protein expression at the single cell level, DNA-barcoded antibody staining was performed (Feature Barcodes). Compatible with the 10x Genomics Next GEM Single Cell 5’ v2 with Feature Barcode chemistry, the TotalSeq™-C Human Universal Cocktail V1.0 (BioLegend #399905) including 137 antibodies recognizing 130 unique cell surface/principal lineage antigens and 7 isotype controls (list of antibodies: https://www.biolegend.com/Files/Images/BioLegend/totalseq/TotalSeq_C_Human_Universal_Co <u>cktail_v1_137_Antibodies_399905_Barcodes.xlsx</u>) was used when applicable. The universal cocktail was designed for the multiomic characterization of peripheral blood-derived immune cells. For optimal characterization of BMA-derived cells, we spiked in the antibodies CD15 (clone W6D3), CD34 (clone 581), and CD117 (clone 104D2) (BioLegend TotalSeq™-C chemistry #C0392, C0054, and C0061, respectively). Briefly, 1 million cells were resuspended into ∼20 uL 1X PBS with 2% FBS. Lyophilized TotalSeq™ panel was equilibrated at room temperature for 5 min, spun down at 10,000 × g for 30 s, resuspended in 27.5 uL 1X PBS with 2% FBS, and spun down again at 20,000 × g for 5 min at 4°C. To the cell suspension, 5 uL of Fc blocking reagent was added (FcX; BioLegend #422301) followed by 5 min incubation. 25 uL of the resuspended TotalSeq™ panel was added, then 1 uL of each of the spike-in antibodies were added for a final volume of ∼50 uL and incubated for 20 min at 4°C. The cells were then washed 3 times (400 × g for 6 min) and counted for quality and quantity assessment using ViaStain AOPI Staining (Nexcelom #CS2-0106-25mL).

### Single cell RNA and Feature Barcode capture

While the focus was on HLA pulldown from cDNA pools barcoded using 10x Genomics 5’ chemistry, we have also conducted the pulldowns and generated data from cDNA captured using 10x Genomics 3’ chemistry, in order to compare both capturing chemistries. Therefore, single cell whole RNA capture and DNA-barcoded cell surface proteins capture (when applicable) were performed using either the 10x Genomics Next GEM Single Cell 3’ v3.1 with Feature Barcode chemistry or the 10x Genomics Next GEM Single Cell 5’ v2 with Feature Barcode chemistry. For the generation of cell-barcoded complimentary DNA (cDNA) molecules, the appropriate 10x Genomics User Guide were followed (document #CG000317_Rev_B for the 3’ chemistry; #CG000330_Rev_A for the 5’ chemistry). Briefly, cells were prepared for capture in 1X PBS with 2% FBS at ∼1,000 cells/uL. About 10,000 cells were submitted for capture using the 10x Chromium Controller. Following reverse transcription and cell barcoding in droplets, emulsions were broken and cDNA was purified using Dynal MyOne SILANE (ThermoFisher Scientific #37002D) followed by a PCR amplification (3’ v3.1 chemistry: 98°C for 3 min; 12 cycles of 98°C for 15 s, 63°C for 20 s, and 72°C for 1 min; 5’ v2 chemistry: 98°C for 45 s; 13 cycles of 98°C for 20 s, 63°C for 30 s, and 72°C for 1 min). Size selection was used to separate the amplified cDNA molecules (> 800 bp) and the DNA from cell surface protein Feature Barcode (< 800 bp) for separate downstream applications, using SPRIselect reagent (Beckman Coulter #B23318). The majority of cDNA molecules in the final product had expected sizes (∼600– 2000 bp).

### Cell surface protein library construction and sequencing

Amplified DNA from cell surface protein Feature Barcodes was used for library construction. Illumina adaptors and TruSeq/Nextera sample indexes (P5, P7, i7 and i5) were added via PCR (98°C for 45 s; 5–7 cycles of 98°C for 20 s, 54°C for 30 s, and 72°C for 20 sec). Amplified DNA libraries were size selected and purified with an expected Agilent Bioanalyzer trace between 200–250 bp. Libraries were sequenced on an Illumina NextSeq2000 sequencer using a pair-ended dual indexing configuration with 28 bp (Read1), 10 bp (i7 Index), 10 bp (i5 Index), 90 bp (Read2) for 3’ capture chemistry, and 26 bp (Read1), 10 bp (i7 Index), 10 bp (i5 Index), 90 bp (Read2) for 5’ capture chemistry, for a sequencing depth of ≥ 5,000 read pairs per cell.

### Gene expression library construction and sequencing

For gene expression library construction, 50 ng (5’ capture chemistry) or between 25–150 ng (3’ capture chemistry) of amplified cDNA were enzymatically fragmented, end-repaired, A-tailed, and ligated to a partial TruSeq adaptor. Additional Illumina adaptors and TruSeq/TruSeq sample indexes (P5, P7, i7 and i5) were added via PCR (98°C for 45 s; 14 cycles of 98°C for 20 s, 54°C for 30 s, and 72°C for 20 sec). Amplified DNA was size selected and purified with an expected size range between 300-600 bp. Libraries were sequenced on an Illumina NextSeq2000 sequencer using a pair-ended dual indexing configuration with 28 bp (Read1), 10 bp (i7 Index), 10 bp (i5 Index), 90 bp (Read2) for 3’ capture chemistry, and 26 bp (Read1), 10 bp (i7 Index), 10 bp (i5 Index), 90 bp (Read2) for 5’ capture chemistry, for a sequencing depth of ≥ 20,000 read pairs per cell.

### Targeted hybridization and magnetic capture of HLA from single cell-barcoded whole transcriptome and long-read sequencing

We designed the scrHLA-typing workflow for in-depth targeted sequencing of HLA genes from a single-cell barcoded whole transcriptome. The 5’-biotinylated hybridization probes were designed to recognize almost all known classical class I and II and non-classical (including pseudogenes) HLA alleles belonging to 40 genes in the IMGT/HLA database^3^, specifically: *HLA-A*, *-B*, *-C*, *-E*, *-F*, *-G*, *-H*, *-J*, *-K*, *-L*, *-T*, *-V*, *-W*, *-Y*, *-DMA*, *-DMB*, *-DOA*, *-DOB*, *-DPA1*, *-DPA2*, *-DPB1*, *-DPB2*, *-DQA1*, *-DQA2*, *-DQB1*, *-DRA*, *-DRB1*, *-DRB2*, *-DRB3*, *-DRB4*, *-DRB5*, *-DRB6*, *-DRB7*, *-DRB8*, *-DRB9*, as well as *HFE*, *MICA*, *MICB*, *TAP1*, and *TAP2*. The workflow can be adapted to any platform capturing single cells and barcoding their transcriptome, including 10x Genomics as described here. The starting material is 10x Genomics-generated full-length cDNA, leftover from what is used to create the Illumina indexed libraries in downstream steps.

#### Optional reamplification

To capture the gene transcripts encoding the HLA molecules, an enrichment was performed using xGen™ Hybridization and Wash v2 kit reagents (IDT #10010351), with modifications. Briefly, 500 ng of barcoded cDNA was needed to start. When not enough cDNA mass was available (< 500 ng), additional cDNA mass was generated by mixing leftover cDNA with KAPA HiFi 2X ReadyMix (Roche #KK2602) and both primers from the 10x cDNA amplification step (10x Genomics #PN-2000089; Suppl. Table S3) at final concentration of 300 nM each, followed by a PCR (98°C for 45 s; 3-to-5 cycles of 98°C for 20 s, 62°C for 20 s, and 72°C for 3 min) and a SPRIselect cleanup (0.6X bead-to-DNA solution ratio).

#### Hybridization and magnetic pulldown

A volume containing 500 ng of cDNA was mixed with 7.5 uL of human Cot1-DNA (supplied in the xGen™ kit) and concentrated on SPRIselect beads using a 1.8X ratio of SPRIselect bead-to-DNA solution suspension. The cDNA was eluted into 19 uL of hybridization mix. In addition to the 2X Hybridization Buffer and the 6.3X Hybridization Enhancer (supplied in the xGen™ kit), the hybridization mix included two blocking oligonucleotides custom-designed to cover the regions at both ends of one of the two strands of a captured cDNA molecule, to block unwanted amplification artifacts. Specifically, where the constant Illumina TruSeq Read1, TSO, and Poly(dT) sequences were located, but also where the highly variable cell barcode and/or unique molecular identifier (CB:UMI) were located. The oligos were modified at their 3’ end by adding a C3 phosphoramidite spacer to prevent them from extending (i.e., acting as primers) in subsequent PCRs (Suppl. Table S3). The blocking oligos’ final concertation in the hybridization mix was 1.6 uM. We also added to the hybridization mix, a mix of biotinylated ∼97-base long probes designed to be highly specific of the HLA transcripts (Suppl. Table S3). The probe mixture final concentration in the hybridization mix was 189.5 nM. The hybridization mix was heated at 95°C for 30 s followed by a rapid ramp down to 65°C which was held for 4 hours allowing the biotinylated probes to bind to the target cDNA. Meanwhile, wash buffers (WB) I, II, and III, and the stringent wash buffer (supplied in the xGen™ kit) were prepared from 10X concentrates with enough volume for two 100 uL washes with WB I, two 100 uL washes with the stringent WB, and one 100 uL wash with each of WB II and III. Streptavidin beads were used to capture the probes bound to their targets. 50 uL of Dynabeads M-270 Streptavidin (ThermoFisher Scientific #65305) were washed two times in a Beads Wash buffer (prepared from the 2X concentrate supplied in the xGen™ kit) and resuspended in the hybridization mix at the end of the 4-hour incubation, immediately followed by an incubation at 65°C for an additional 45 min with frequent gentle hand mixing every 10–15 min. At the end of the incubation, the beads were washed on a magnetic rack one time with WB I pre-heated at 65°C and two times with stringent WB pre-heated at 65°C followed by a series of one-time washes with WB I, II, and III, at room temperature. At the end of the last wash, the beads were removed from the magnet and resuspended in EB buffer (Qiagen #19086).

#### Amplification of hybridization product

Following hybridization capture of HLA molecules, an on-bead PCR was performed to amplify captured molecules. The product (beads + captured cDNA) was mixed with KAPA HiFi 2X ReadyMix (Roche #KK2602) and both primers from the 10x cDNA amplification step (10x Genomics #PN-2000089; Suppl. Table S3) at final concentration of 300 nM each, followed by a PCR (98°C for 45 s; ∼10–14 cycles of 98°C for 20 s, 62°C for 20 s, and 72°C for 3 min), and a SPRIselect cleanup (0.8X bead-to-DNA solution ratio).

#### TSO artifact removal

A significant fraction of the cDNA product from a 10x Genomics single cell cDNA preparation (3’ or 5’ chemistry) may contain a template switch oligo (TSO) priming artifact (i.e., TSO sequence at both ends of the cDNA molecule) instead of the correct structure. TSO artifact depletion is recommended for long-read sequencing on platforms such as Pacific Biosciences. Following hybridization capture and PCR-amplification of the pulldown product, TSO artifact depletion was performed^36^ by first mixing the amplified pulldown product with KAPA HIFI 2X Uracil+ (Roche #KK2802), a biotinylated forward primer with two uracil additions at the 5’ end, and a reverse primer. Final primer concentration was 500 nM. The biotinylated uracil primer captured the constant region on the opposite end of where the CB:UMI and TSO sequences are located on the cDNA molecule while the regular primer captured the constant region that encompassed the CB:UMI and TSO sequences (Suppl. Table S3). The mix was amplified by PCR (98°C for 45 s; ∼4 [if cDNA input ∼250 ng] to 6 [if cDNA input ∼25 ng] cycles of 98°C for 20 s, 65°C for 30 s, and 72°C for 4 min) followed by a SPRIselect cleanup (0.8X bead-to-DNA solution ratio). cDNA molecules with the correct structure were then captured and purified using 10 uL (100 ug) Dynabeads™ kilobaseBINDER™ streptavidin beads (ThermoFisher #60101) in 40 uL of provided Binding Solution, incubated 15 min at room temperature, washed on magnet 2 times by provided Washing Solution and 1 time by distilled water, and finally reconstituted in 40 uL EB. After streptavidin binding, 2 uL of USER enzyme (NEB #M5505L) was added to the bead/cDNA mixture and incubated at 37°C for 30 min to uncouple the cDNA from the beads. After magnetic separation, the supernatant containing the cDNA was purified by a SPRIselect cleanup (0.8X bead-to-DNA solution ratio) and quantified.

#### TSO-depleted reamplification for blunt-end product

The sample was reamplified after TSO depletion, to get back regular blunt ends on the product, by mixing it with KAPA HiFi 2X ReadyMix (Roche #KK2602) and both primers from the 10x cDNA amplification step (10x Genomics #PN-2000089; Suppl. Table S3) at final concentration of 300 nM each, followed by a PCR (98°C for 45 s; 4 cycles [if ≤ 125 ng input cDNA], 3 cycles [if 125–500 ng input cDNA], or 2 cycles [if ≥ 500 ng input cDNA] of 98°C for 20 s, 62°C for 20 s, and 72°C for 3 min) and a SPRIselect cleanup (0.8X bead-to-DNA solution ratio).

#### Constructing PacBio SMRTbells

Our standard long-read sequencing approach was to multiplex up to 4 hybridization capture products on a PacBio ‘single molecule real-time’ (SMRT) Cell 8M, using the ‘Iso-Seq’ method. Following TSO-depletion of the pulldown captured cDNA, the product was DNA-repaired, A-tailed, SMRTbell overhang adaptor-ligated (each sample with one of the barcoded Overhang Adaptors 8A/8B [PacBio #101-628-400/500] when multiplexing samples on a single SMRT Cell), nuclease-treated, and SMRTbell-cleanup purified (1X bead-to-DNA solution ratio) using the PacBio SMRTbell Prep kit 3.0 (PacBio #102-141-700), as per manufacturer’s protocol (PacBio Procedure & Checklist #102-359-000 REV03 DEC2023). Libraries were sequenced on a PacBio Sequel IIe with 2-hour pre-extension time and 30-hour movie time.

### Augmenting long-read throughput by multiplexed arrays isoform sequencing (MAS-seq)

When augmenting throughput by MAS-seq was desired, the following intervention to the hybridization pulldown workflow was performed. Following TSO-depletion and right before the blunt-end reamplification, the TSO-depleted product was brought through the MAS-PCR step of the PacBio Procedure & Checklist for MAS-seq library prep (PacBio #102-678-600 REV03 MAR2023), inspired by Al’Kafaji et al.’s design^36^. Briefly, after TSO depletion, 50 ng of the TSO-depleted product in 45 uL of buffer was mixed with 125 uL of nuclease-free water and 212.5 uL of MAS-PCR mix (PacBio #102-692-800) or KAPA HIFI 2X Uracil+ (Roche #KK2802), was split into 16 tubes (22.5 uL per tube) to each of which 2.5 uL of MAS primers premixes ‘A’ through ‘P’ (provided in PacBio’s MAS-seq kit #102-659-600) was added, and PCR-amplified (98°C for 3 min; 9 cycles of 98°C for 20 s, 68°C for 30 s, and 72°C for 4 min). Products from the 16 reactions were pooled and cleaned up using either SMRTbell cleanup beads (PacBio #102-158-300) or ProNex Size Selection Chemistry beads (Promega #NG103B) using a 1.5X bead-to-DNA solution ratio. The rest of the USER enzyme (NEB #M5505L) digestion, ligation of pooled product into arrays sealed on both their ends with MAS adaptors, DNA damage repair and nuclease treatment of the arrays, was conducted according to manufacturer’s protocol (PacBio #102-678-600 REV03 MAR2023). Arrays were sequenced on a PacBio Sequel IIe with 2-hour pre-extension time and 30-hour movie time.

### Targeted hybridization and magnetic capture of specific mutated transcripts from single cell-barcoded whole transcriptome and long-read sequencing for gene-fusion detection or short-read sequencing for SNV detection

We designed a highly adaptable workflow for targeted hybridization capture and amplification of gene transcripts with known mutations or fusions from the whole amplified transcript pool (cDNA). For patients ‘AML1’, ‘AML4’ and ‘AML6’, mutations of the single nucleotide variants (SNVs) type and/or gene fusions were identified elsewhere, through clinically validated commercial assays. Once identified, 5’-biotinylated hybridization oligonucleotide probes appropriately specific to the gene(s) carrying the mutation(s) were designed (Suppl. Table S3). If destined for short-read sequencing, we additionally designed reverse PCR-primer oligonucleotide(s) positioned a few nucleotides downstream each SNV (Extended Data Fig. 1a). The fusion-specific and long-read sequencing experimental protocol was the same as for HLA targeted capture (see above) except that the biotinylated capture probes added to the hybridization mix were specific to the fusion breakpoints (Extended Data Fig. 12a). The SNV-specific and short-read sequencing experimental protocol followed the same first steps as for the HLA targeted capture (see above), with a starting material at 500 ng of cell-barcoded full-length cDNA (here, generated by the 10x Genomics platform), and the ‘optional reamplification’ and ‘hybridization and magnetic pulldown’ steps. Following hybridization capture, an on-bead targeted PCR was performed to amplify regions that included the mutations of interest. The forward primer universally recognized 10x Genomics-generated cDNA molecules and started at the CB:UMI location, while the reverse primer(s) were designed to specifically bind to a region 40∼70 bp downstream the mutation on the captured transcripts. The PCR mix consisted of the KAPA HiFi 2X ReadyMix (Roche #KK2602), the captured cDNA molecules on their beads, a universal 10x Genomics compatible forward primer, and as many custom-designed reverse primers as there were mutations to resolve. Each primer was at a final concentration of 500 nM. A PCR was then carried out (98°C for 45 s; 14–18 cycles of 98°C for 20 s, 62°C for 20 s, and 72°C for 3 min), followed by a SPRIselect cleanup step while carefully choosing a size selection ratio compatible with the expected size of the smallest amplicon in the PCR product. As a quality control measure, the product size was assessed and the number of peaks in the trace profile corresponded to the number of expected amplicons in the PCR product (Extended Data Fig. 1b). Following targeted PCR, the amplicons were end-repaired, A-tailed, and adaptor-ligated using the NEBNext® Ultra™ II End Repair/dA-Tailing and Ligation Module reagents (NEB #E7546S/L and #E7595S/L). 50 uL of captured and target-amplified cDNA was mixed with 7 uL of End Prep buffer and 3 uL of End Prep enzyme, and incubated 30 min at 20°C then 30 min at 65°C. After incubation, 30 uL of Ligation Mix, 1 uL of Ligation Enhancer, and 7 uL of 10x Adaptor Oligo (10x Genomics #PN-2000094) were added to the previous mix and incubated 20 min at 20°C. A SPRIselect cleanup followed, and the eluted product underwent an indexing PCR (98°C for 45 s; 7 cycles of 98°C for 20 s, 54°C for 30 s, and 72°C for 3 min) to add Illumina compatible TruSeq/TruSeq P5, P7, i7, and i5 adaptors using the 10x Genomics Dual Index Plate TT Set A (10x Genomics #PN-3000431) mixed with the KAPA HiFi 2X ReadyMix. Libraries were sequenced on an Illumina NextSeq2000 sequencer using a pair-ended dual indexing configuration with 26 bp (Read1), 10 bp (i7 Index), 10 bp (i5 Index), 90 bp (Read2), for a sequencing depth between 500–2,000 read pairs per cell.

### Single cell gene expression and cell surface protein data processing with Seurat

Illumina BCL files were demultiplexed and converted to FASTQ format using ‘bcl2fastq’ version 2.20.0 (Illumina, Inc.) through the cellranger-mkfastq function (10x Genomics, Inc.). Resulting FASTQ files were processed with ‘cellranger’ (version 6.0.2) to identify and count Feature Barcodes, and to align to the hg38 reference genome and count Gene Expression data. An output filtered feature barcode matrix file was then read on R^54^ (version 4.2.0) and converted into a ‘Seurat’ object^55^ (version 5.0.1). Initial quality control was used to subset out cells with too many (>50,000) or too few (<1,000) RNA counts and too many mitochondrial RNA counts (>20% of a cell’s total RNA counts). Counts were then normalized assuming proportionality to the sum of counts of every feature per CB (‘LogNormalize’ method, scale factor: 10,000), variable features were selected (‘vst’ selection method), features were centered and scaled (‘ScaleData()’ function), principal component analysis on the variable features was run, and UMAP low dimensional embedding^56^ and clustering using k-nearest neighbor algorithm was performed on data.

### Automatic cell type labeling using viewmastR

The cell types were automatically labeled using the ‘viewmastR’ package^31,57^ (version 0.2.1), a machine-learning platform performing unsupervised classification of single cells between the query dataset and a well-annotated reference dataset using multinomial logistic regression, relying on the top 5,000 genes commonly variable across the datasets. In our study of BMMCs from bone marrow aspirates, we used a previously annotated reference dataset of healthy marrow and peripheral blood mononuclear cells^32^.

### Genetic demultiplexing of allogeneic entities with souporcell

To perform variant calling and distinguish donor vs. recipient cells of bone marrow samples post-alloHCT, first the raw sequence data aligned to hg38 files (the BAM files) output from cellranger were merged using ‘mergebams’ (version 0.3), a custom script that preserves sample tags onto CBs^58^. Then, ‘souporcell’^30^ (version ≥ 2.0; ‘singularity’ version 3.5.3) was used on the merged BAM files as input, the cellranger output sample filtered cell barcodes as the barcode file, the hg38 reference sequence as the reference file, a variants file (VCF format) from the 1000Genomes project^59^ as the common variants file, an integer number ‘k’ as the number of expected genotypes in the sample, and invoking the ‘ignore’ and ‘skip_remap’ options. Sparse mixture model output from souporcell was log-normalized and colored by the genotype assignment.

### IsoSeq computational processing of long-read HLA pulldown sequencing data

Error-corrected sequencing reads were generated by the Circular Consensus Sequencing (CCS) workflow, on-board the PacBio Sequel IIe instrument (Pacific Biosciences of California, Inc.), generating a high fidelity ‘HiFi’ reads BAM file. When HiFi reads were concatenated (i.e., from MAS-seq), an additional segmentation step was performed by providing the list of MAS adaptors and running the ‘split’ program in the PacBio concatenated read splitter ‘Skerà package. HiFi or segmented HiFi reads were then processed in the ‘IsoSeq’ suite of computational tools (version 3.8.2) by following those programs in order: 1) ‘limà (version 2.7.1) was used to remove sequencing primers, CTACACGACGCTCTTCCGATCT (5p) and GTACTCTGCGTTGATACCACTGCTT (3p) for the 10x 5’ chemistry reads, and AAGCAGTGGTATCAACGCAGAGTACATGGG (5p) and AGATCGGAAGAGCGTCGTGTAG (3p) for the 10x 3’ chemistry reads, 2) ‘tag’ was used to separate the UMI and CB from the rest of the transcript using the ‘16B-10U-T’ design (10x 5’ chemistry) or the ‘T-12U-16B’ design (10x 3’ chemistry), 3) ‘refinè was used to trim the polyA sequence and remove unintended concatemers (if not already done by Skera), and 4) ‘correct’ was used to identify cell barcode errors and correct them against the 10x 5’ chemistry CB whitelist or the 10x 3’ chemistry reverse complemented CB whitelist. All of the IsoSeq pipeline steps were used with default settings.

### scrHLAtag to align long-read HLA pulldown sequencing data to the IMGT/HLA reference

We developed ‘scrHLAtag’ (version 0.1.7), a command line tool written in Rust for processing full-length single cell-barcoded HLA transcript-enriched PacBio long-read sequencing data from 10x Genomics. We designed scrHLAtag to use ‘minimap2’ (version 2.24), primarily designed for fast and high-accurate alignment of long-reads^60^, to align sequences to the mRNA/cDNA reference library (exons only) and to the genomic reference library (exons, introns and/or UTRs) of all known HLA alleles available from the IMGT/HLA database^3^ (version: 3.60.0 – 2025-04).

Because scrHLA-typing is based on single cell RNA-seq data, HLA alleles differing in non-coding regions cannot be resolved by our assay. Therefore, only up to 3-field HLA allele names were used. HLA alleles which only varied in the 4th field from each other (e.g., *A*\*30:02:01:01, *A*\*30:02:01:02, *A*\*30:02:01:03, etc.) were reduced to one 3-field allele in the references, where the sequence of only the corresponding XX:XX:XX:01 allele was retained. The 42,579 cDNA alleles and the 24,355 genomic alleles known from IMGT/HLA version 3.60.0 became 31,141 and 16,916 alleles (respectively) after reduction to 3 fields.

The IsoSeq ‘corrected’ BAM files were used as input to scrHLAtag. The output of scrHLAtag included BAM files aligned to the IMGT/HLA transcriptome and/or genome indexed and sorted by read name (‘SAMtools’; version 1.16.1), file(s) summarizing read counts per CB:UMI, FASTA references used by minimap2, and molecule information files with each line representing one read count and containing the sequencing read with its associated CB and UMI, the allele to which it was mapped, and other minimap2 alignment metrics.

Running unsupervised (i.e., without providing a predetermined list of HLA alleles), scrHLAtag aligns the ‘corrected’ BAM to the entire IMGT/HLA reference library. This typically yields count files with a few thousands unique HLA alleles mapped to the reference (between 4853 and 7828 unique alleles in our study; Suppl. Table S1), a number considered too high given that the target genes are 40, but not surprising given the high level of homology between the tens of thousands of alleles. scrHLAtag can also run ‘supervised’ or ‘guided’ if provided with a short list of HLA alleles. This list typically contains names of the top HLA candidates progressively refined via scrHLAmatrix (see below) after multiple scrHLAtag–scrHLAmatrix iterations using the same ‘corrected’ BAM alongside the newly refined list as input. The convergence of the HLA candidate list to the putative HLA genotypes of each allogeneic entity in a given sample is known when such list stops updating after additional iterations.

### scrHLAmatrix to predict HLA genotypes, perform deduplication, and summarize counts from scrHLAtag output

To analyze scrHLAtag output, we developed ‘scrHLAmatrix’ (version 0.18.1), a package written in R to process scrHLAtag molecule information read count files to predict HLA genotypes in single cells, remove PCR duplicates, and summarize counts into Seurat-compatible^55^ matrices (Fig. 1b).

#### Filtering by read quality

Best quality alignment reads in molecule information count files were identified and retained. scrHLAtag collected from minimap2 a combination of metrics including the ‘s1’ (chaining score), ‘AS’ (dynamic programming alignment score), ‘NM’ (total number of mismatches and gaps in the alignment), and ‘de’ (gap-compressed per-base sequence divergence) tags. Assuming imperfections during alignment, we hypothesized the scores in these tags would be unfavorable if a query sequence was mapped to the wrong reference sequence. We exploited the availability of ‘Donor’ vs. ‘Recipient’ labeling of sequences by souporcell as ‘ground truth’ to compare the score values in correct vs. wrong mappings. Score thresholds of good quality mappings were set where the sum of True Positive Rates (donor or recipient mappings correctly validated by souporcell) and True Negative Rates (donor or recipient mismappings as predicted by scores, correctly refuted by souporcell) was maximized at each HLA locus, for each of the tags (Suppl. Fig S1a). Across the loci and their alleles, on average, thresholds for ‘s1’ and ‘AS’ were such that good quality mappings had a percentage passing score of 81.8 and 80.7, respectively, from the maximum scores of each locus. For ‘NM’ and ‘de’, good quality mappings had scores at or below 21 and 0.004, respectively, on average across the loci (Suppl. Fig S1a). Thus, we set default ‘s1’ and ‘AS’ percent passing scores, and default ‘NM’ and ‘de’ thresholds in our scrHLAmatrix functions at 80, 80, 15 (slightly more stringent), and 0.01 (slightly less stringent), respectively (examples in Suppl. Fig S1b).

#### Classifying cells based on HLA genotype patterns

With the presence of chimeric entities within the captured single cells, we hypothesized the distribution of identified HLA alleles among the cells would reveal patterns, mostly driven by the recipient-donor mismatched alleles. To visualize these patterns, molecule information files were first organized into ‘HLA × CB’ count matrices. With the option to match the CBs of the count matrix with those of the associated Seurat object (allowing the analysis of previously quality controlled CBs), the counts of each allele were normalized assuming proportionality to the sum of counts of every HLA allele per CB (normalization by size factor) and log-transformed. Next, a principal component analysis (PCA) was performed to emphasize the patterns. By default, up to the top 50 PCs were retained for downstream clustering and UMAP (uniform manifold approximation and projection) analyses. Increasing or decreasing this number according to the scree plot or ‘elbow’ plot shape, was optional to retain the top PCs accounting for most of the variation and discard the bottom ‘noisy’ PCs. To classify cells into groups with common HLA genotype expression patterns, a graph-based clustering approach was applied. Algorithm choices included a community structure detection method: the Leiden algorithm^61^ (‘leiden’: version 0.4.3), a density-based method: DBSCAN^62^ (‘dbscan’: version 1.1.10), a connectivity-based method: hierarchical clustering (agglomerative), a centroid-based method: k-means (both ‘hclust’ and ‘kmeans’ provided by the ‘stats’ package part of R^54^, version 4.2.0), and a distribution-based method: Gaussian Mixture Model (provided by the ‘mclust’ package^63^, version 6.0.0). All clustering algorithms were applied to PCA space, except DBSCAN, known for its degraded performance on multidimensional data^64^. Instead, we performed non-linear manifold-aware dimension reduction of the retained PCs with UMAP^65^ (‘uwot’: version 0.2.2) down to 2 dimensions, onto which we applied ‘dbscan’. We further tuned UMAP parameters to generate denser clumps and cleaner separations between clusters for better DBSCAN performance—specifically, setting ‘min_dist’ and ‘repulsion_strength’ to the low values of 0.001. The number of genotype classes could be fixed a priori and in this study was equal to 2 (‘k = 2’) representing the donor and recipient entities in our samples, although that number could be greater than 2 (see below regarding in-silico mixing ≥ 3 chimeric entities). We did not set assumptions on which algorithm had the best ability to correctly classify cells into the different chimeric entities. Instead, we extracted clustering ‘*consensus*’ by grouping the CBs into the same cluster if they agreed on membership in a majority of clustering algorithms clusters (low confidence classifications were set as ‘not assigned’). The consensus algorithm’s performance in correctly predicting the chimeric entities depended on the number of donor-vs.-recipient mismatched alleles among total alleles, with greater proportion of mismatches leading to classification accuracy close to 100% (Extended Data Fig. 14; and Extended Data Fig. 8 to see examples of HLA pattern-based classification vs. souporcell classification in a patient with 3 samples at various levels of chimerism).

#### Predicting HLA genotypes

A ‘per-single-cell’ prediction algorithm was performed, consisting first of summarizing counts into ‘HLA × CB’ matrices for each identified cluster of cells, separately. Then, an account of the top 2 most numerous alleles per loci and per CB was performed. The ‘top 2 alleles’ was based on the assumption that a cell would normally have at most 2 alleles per locus. Another assumption was that correctly aligned alleles would be more abundant than misaligned or slightly misaligned alleles, thus more likely to be retained in a per-cell tally. Ties where ≥ 3 alleles were equally abundant or where ≥ 2 alleles were equally abundant at the 2nd place, were resolved by eliminating tied alleles due to undecidability. The resulting genotypes per loci per CB, either apparent heterozygous (i.e., ‘Allele1_Allele2’) or apparent homozygous (i.e., ‘Allele1_blank’) were then ranked from most to least abundant for each chimeric cluster. The highest-ranking genotypes cumulatively accounting for a majority of total counts for a locus (by default 85% of the total) were retained in the final list of candidate alleles. When scrHLAtag runs returned several thousand uniquely mapped alleles (most commonly during the 1st ‘unsupervised’ runs; examples in Suppl. Table S1), the ‘HLA × CB’ count matrices would become too sparse, and the per-single-cell prediction algorithm would no longer yield accurate genotype predictions. By default, when the number of unique HLA alleles was > 3000, a simpler ‘pseudo-bulk’ prediction algorithm was used. The algorithm handled the molecule information count files as bulk data frames. For each identified cluster of cells, the highest-ranking mapped alleles in terms of bulk counts, cumulatively accounting for a majority of total counts for a locus (by default 85% of the total) were retained in the final list of candidate alleles. The list of candidate alleles was then ready to be supplied to the next run of scrHLAtag.

Note that among the 31,141 reference entries of the mRNA/cDNA 3-field reduced IMGT/HLA library, 6 of them had an extra 24 to 99 validated nucleotides downstream the 3’ furthest most common sequencing starting position in the reference library (Extended Data Fig. 15). The alleles were: *DPA1*\*02:38Q, *A*\*03:437Q, *B*\*13:123Q, *C*\*02:205Q, *C*\*04:09L (previously named *C*\*04:09N), and *C*\*04:61N, and considered rare in the populations. When the PacBio long-read RNA sequences—often longer than typically available reference sequence length—of the more common alleles *DPA1*\*02:02:02, *A*\*03:01:01, *B*\*13:02:01, *C*\*02:02:02, *C*\*04:01:01, and again *C*\*04:01:01 were aligned to the reference library, minimap2 of scrHLAtag preferentially matched them, respectively, to those erroneous references, despite incurring a 1 to 2 nucleotide(s) mismatch penalty (note: between *C*\*04:09L and *C*\*04:61N, a *C*\*04:01:01 sequence would always preferentially misalign to the former, because it would incur fewer penalties vs. the latter; Extended Data Fig. 15d,e). To mitigate this, an option was made available to adjust the final list of candidate alleles accounting for those 6 known misalignments.

#### Post-alignment PCR-deduplication

Algorithms to remove PCR duplicates based on a ‘consensus quality value’ guided approach to generate one consensus sequence per founder (such as the ‘Dedup’ program in IsoSeq) were avoided. Because mismatches and shifts ‘naturally’ occur in HLA, we suspected these types of deduplication might create consensus amalgams not fully aligning to any allele, thus not appropriate for HLA sequencing where the difference between alleles is sometimes a single nucleotide mismatch. Moreover, because of the high degree of nucleotide homology between alleles of the same HLA gene or different genes but of the same HLA class, the template switching PCR artifact (i.e., molecular swap, noted in some high quality read clusters with the same CB:UMI; Extended Data Fig. 3) was prevalent. To account for this and also to avoid ‘chimeric’ consensuses, an enumeration-based, post-alignment (with minimap2 of scrHLAtag) deduplication strategy was adopted, where among reads belonging to the same CB:UMI, the allele with the most counts prevailed. When ties were present with the most numerous aligned allele, the entire CB:UMI was dropped due to undecidability.

#### Summarizing counts in single cells in Seurat-compatible matrices

After running scrHLAtag for the final time (i.e., with the most curated list of candidate alleles), reads were filtered by minimap2 quality read tags (see above) and PCR-deduplicated (see above). Additional options for read filtering were available (Extended Data Fig. 9a). These included resolving donor-vs.-recipient genotype conflict (assuming a cell cannot have both recipient and donor-origin HLA allele) by keeping either donor-specific or recipient specific HLA-associated UMIs (when known), based on their count difference per cell. Another option was resolving per-HLA genotype conflicts, assuming each cell had a maximum of 2 genotypes per HLA gene and keeping those with the most counts. A third option was correcting linkage disequilibrium (LD) in the HLA-DR subregion, assuming well-established strong LD in the DR1 haplotype (*DRB1*\*01 and *10 in LD with *DRB6* and *DRB9*), the DR51 haplotype (*DRB1*\*15 and *16 in LD with *DRB6*, *DRB5*, and *DRB9*), the DR52 haplotype (*DRB1*\*03, *11, *12, *13, and *14 in LD with *DRB2*, *DRB3*, and *DRB9*), the DR8 haplotype (*DRB1*\*08 in LD with *DRB9*), and the DR53 haplotype (*DRB1*\*04, *07, and *09 in LD with *DRB7*, *DRB8*, *DRB4*, and *DRB9*), and removing reads associated with genes that are in conflict with the LD rules of the abovementioned DRB1 allele families. In practice, these additional options allowed the fine-graining of the reads count files without dramatically changing the counts (Extended Data Fig. 9c). Finally, the reads were summarized in ‘HLA × CB’ sparce numeric matrices.

### Assessing mismatch patterns in HLA sequencing reads across single cells

Bioconductor tools were used to summarize and visualize mismatch patterns in HLA sequencing reads across single cells. The 3-field reduced HLA mRNA reference from scrHLAtag was forged into a ‘BSgenomè class object^66^. Then, the long-read HLA-aligned BAM files from each CD34-sorted and unsorted bone marrow specimens of each sample were first merged then subset into distinct groups of interest by CB (i.e., healthy donor, healthy recipient, or malignant recipient cells) using ‘mergbamsR’ version 0.0.5, and R package we designed for the purpose^67^. Finally, using ‘Rsamtools’ version 2.14.0 from Bioconductor^68^, alignment statistics were extracted and summarized.

### In-silico mixes with 3 or more chimeric entities to test the HLA genotype-based cell classifier

In-silico mixes were necessary because samples naturally consisting of 3 or more chimeric entities were not available in this study (the closest clinically available candidates would have been a sample from a patient post-double cord blood transplant or a from a patient with a second or third transplant; however, even in those cases, one donor usually prevails^69^). CBs from the Seurat objects of AML4_d90, AML8, AML6, and AML1 were extracted based on whether their labels were ‘Donor’ or ‘Recipient’ (by souporcell). The ‘corrected’ BAM files (output of PacBio’s IsoSeq *Correct* program) were subset using SAMtools to only retain the reads associated the extracted CBs. Using mergebams, mixtures of 3, 4, 5, and 6 chimeric entities were created, making sure to tag sample labels to their corresponding CBs. Post-merge BAMs were then input to scrHLAtag, in an unsupervised way initially, and subsequently into multiple rounds of scrHLAtag guided by a gradually converging list of HLA candidates predicted by scrHLAmatrix. Given a value ‘k’ of ‘3’, ‘4’, ‘5’, or ‘6’ respectively corresponding to each of the 4 in-silico mixes, the scrHLAmatrix classifier grouped cells into 3, 4, 5, or 6 clusters based on HLA expression patterns. The classification accuracy, calculated as the percentage of cells within a given classification corresponding to the same majority label (known a priori), ranged from 98.2% to 100% (Extended Data Fig. 4).

### Mutational short-read computational data processing

Illumina BCL files were demultiplexed and converted to FASTQ format using bcl2fastq through the cellranger-mkfastq function, as the mutational targeted sequencing and hybridization capture workflow was adapted to the 10x Genomics workflow. Resulting FASTQ files were then processed with ‘mütCaller’ (version 0.7.0), a command line tool written in Rust we developed to call mutations in sequencing data^70^. To align sequences to the hg38 reference genome, mütCaller used minimap2 by default when provided with unaligned FASTQs. The other option was ‘STAR’^71^ (version 2.7.2a). mütCaller then counted CB:UMIs with reference nucleotides and with alternative nucleotides in the alignment BAM files based on a list of a priori reference vs. alternative (or query) nucleotides at a given chromosomal location. A given UMI for a given CB was considered ‘reference’ genotype if reference counts for the CB+UMI combination was greater than alternative counts, and vice versa. Then ‘reference’ and/or ‘alternative’ UMIs for a given CB were summarized and such CB was called as expressing the ‘reference’ (i.e., wildtype) transcript, the ‘alternative’ (i.e., mutant) transcript, or both (i.e., biallelic).

### Fusion-gene long-read computational data processing

First, the generation of ‘HiFi’ reads BAM files from CCS reads was performed. Then the following IsoSeq pipeline steps (with default settings) were performed in the following order: ‘limà, ‘tag’, ‘refinè, ‘correct’, and ‘groupdedup’. The resulting PCR-duplicate-corrected BAM files were aligned to the hg38 reference genome as recommended by PacBio^72^ using ‘pbmm2’ version 1.13.1 (Pacific Biosciences of California, Inc.) with program ‘align’ and options ‘--sort’ and ‘--preset ISOSEQ’. Gene fusions were then counted and summarized into Paired-End BED files using ‘pbfusion’ version 0.5.0 (Pacific Biosciences of California, Inc.) with program ‘discover’ using a serialized GTF (general transfer format) file and the option ‘--min-fusion-quality MEDIUM’.

## Data and code availability

Normalized count matrices and metadata are provided for review. All code is available on https://github.com/furlan-lab/scrHLA_manuscript. Full data including all raw sequence have been deposited in NCBI’s Gene Expression Omnibus^73^ and are accessible through GEO Series accession numbers GSE305305, GSE305306, and GSE305307 (https://www.ncbi.nlm.nih.gov/geo/query/acc.cgi?acc=GSE305305, https://www.ncbi.nlm.nih.gov/geo/query/acc.cgi?acc=GSE305306, and https://www.ncbi.nlm.nih.gov/geo/query/acc.cgi?acc=GSE305307).

## ACKNOWLEDGEMENTS

We thank Dr. Luca Vago at the Ospedale San Raffaele for his helpful comments, the Fred Hutchinson Scientific Computing Group and Genomics Core as well as the Seattle Children’s Tumor Bank and the SCH-B802 study team for providing the resources to carry out this work. This study was supported by multiple sources of funding, including the National Institutes of Health/National Cancer Institute (NIH/NCI) grant R01-CA-289886 to S.N.F, S10-OD-020069 and S10-OD-028685 to the Fred Hutch/University of Washington/Seattle Children’s Cancer Center Consortium NCI Cancer Center Support Grant P30. The Hartwell Foundation provided support for a portion of this work via an Individual Biomedical Research Award to S.N.F.

## AUTHOR CONTRIBUTIONS

S.B.K. and S.N.F. designed research. S.B.K., E.F., and S.S.B. processed samples and performed experiments. S.B.K., J.G.U., and S.N.F. performed and/or provided input on data processing, coding, and analysis. R.G.G., M.S.T., C.A.J., J.S., A.N.G., S.R.R., M.B., and S.M. provided access to patient samples and assisted with processing. S.B.K. and S.N.F. wrote the manuscript with input from all authors.

## EXTENDED DATA FIGURES AND LEGENDS

**Extended Data Fig. 1.**
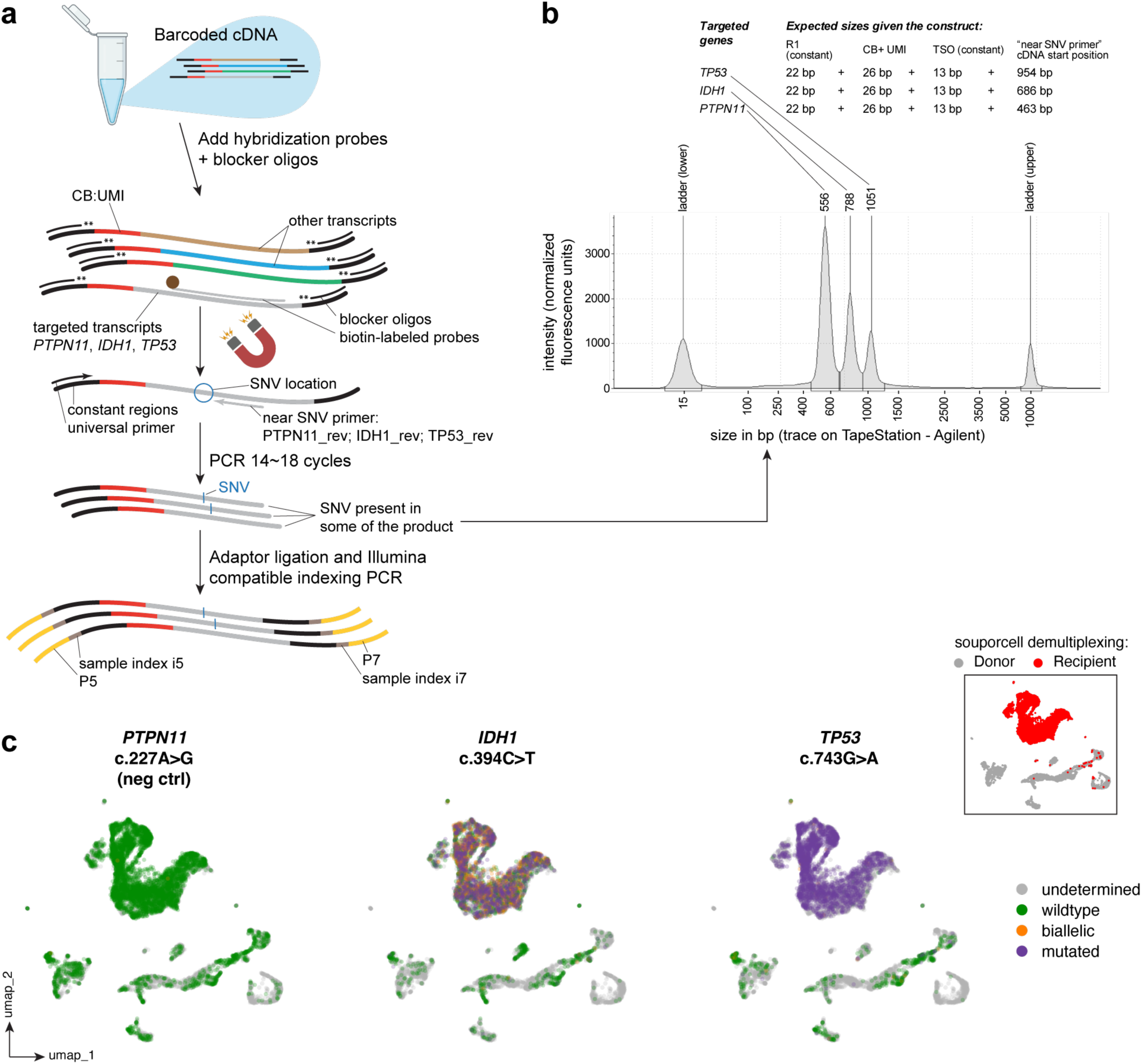
Experimental design and output of targeted gene mutation capture in AML1. **a**, Starting product is single-cell barcoded cDNA pools (from a capture platform like 10x Genomics). To that, custom-designed biotinylated hybridization probes and blocking oligos are added. Targeted transcripts are enriched magnetically (using streptavidin beads); PCR-amplified using a universal ‘forward’ primer specific to the constant region upstream of the barcode and ‘reverse’ primer(s) (as many as there are targeted genes) specific to a region near the known single nucleotide variants (SNVs) on the targeted gene; and prepped for Illumina-compatible short-read sequencing. **b**, Representative DNA trace (TapeStation; Agilent Technologies, Inc., Santa Clara, CA, USA) of actual PCR product after hybridization capture and amplification of 3 target transcripts: *PTPN11*, *IDH1*, and *TP53*, along with the ‘expected’ sizes of those products given the locations of the forward and reverse primers. **c**, Using ‘mütCaller’ (a pipeline we designed for identifying mutations in single cell data) to identify two known mutations in the *IDH1* and *TP53* genes (clinically assessed, in patient file) and a third arbitrary mutation on the *PTPN11* gene not known to be present within the patient (negative control), mutated *IDH1* and *TP53* transcripts were found associated with cell barcodes later identified as leukemia cells. By counting mutations on UMIs per each cell barcode (CB), cells were either expressing mutated transcripts, a combination of mutated and wildtype transcripts (biallelic), wildtype transcripts, or expression of targeted transcript was not detected (undetermined). Classification of cells by genetic demultiplexing (souporcell) into recipient and donor is also shown for reference.

**Extended Data Fig. 2.**
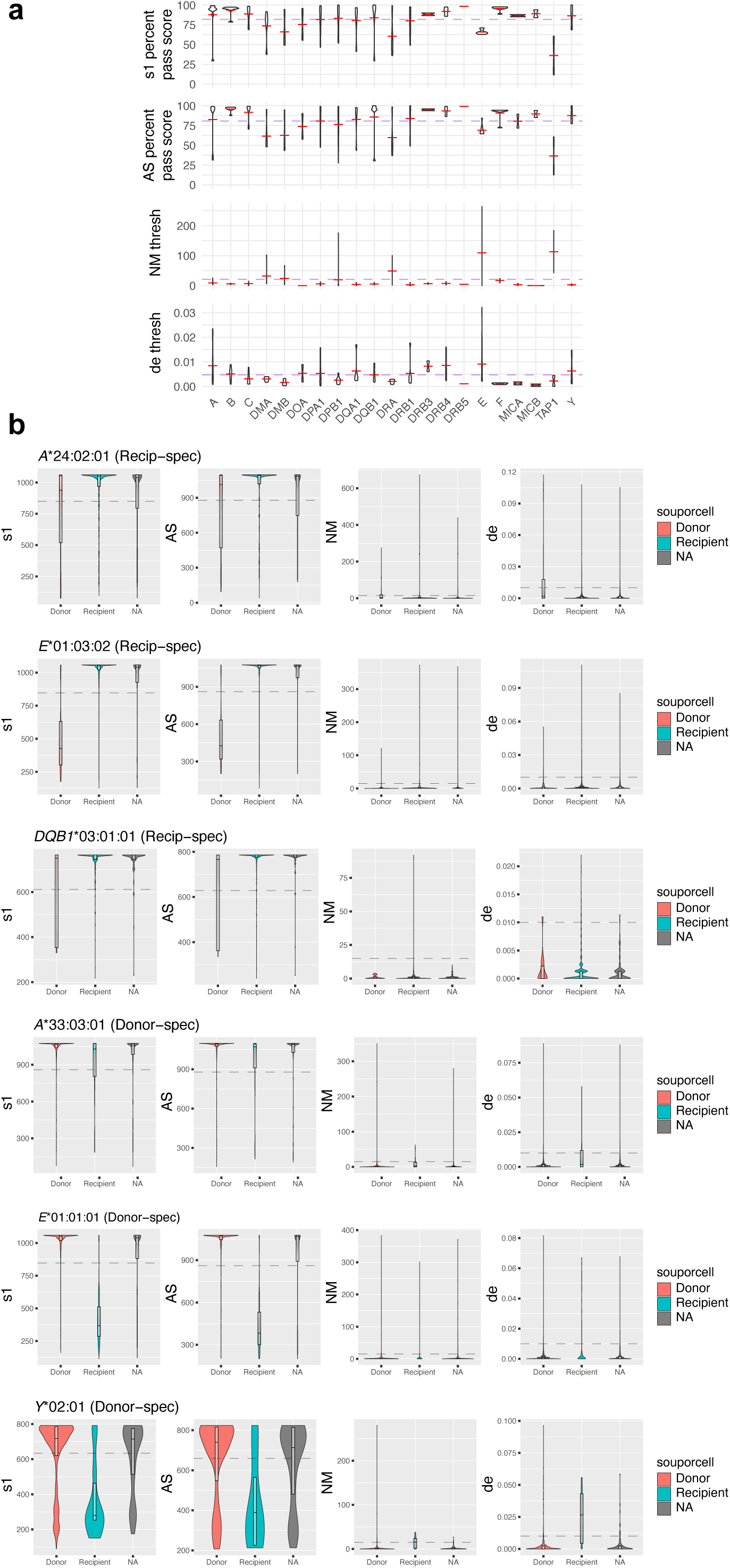
Identifying best quality reads according to minimap2 tags. **a**, Score thresholds of good quality mappings according to the ‘s1’ (chaining score), ‘AS’ (dynamic programming alignment score), ‘NM’ (total number of mismatches and gaps in the alignment), and ‘de’ (gap-compressed per-base sequence divergence) tags, where the sum of True Positive Rates (donor or recipient mappings correctly validated by souporcell) and True Negative Rates (donor or recipient mismappings as predicted by scores, correctly refuted by souporcell) was maximized at each HLA locus, for each of the tags (purple dotted line represents the mean threshold of each gene). **b**, Examples taken from several alleles (with an indication of whether they were recipient or donor-specific) and how the scores distribute in cell barcodes associated with recipient cells, donor cells, or not appearing (NA) in the corresponding Cell-Dataset Object column names.

**Extended Data Fig. 3.**
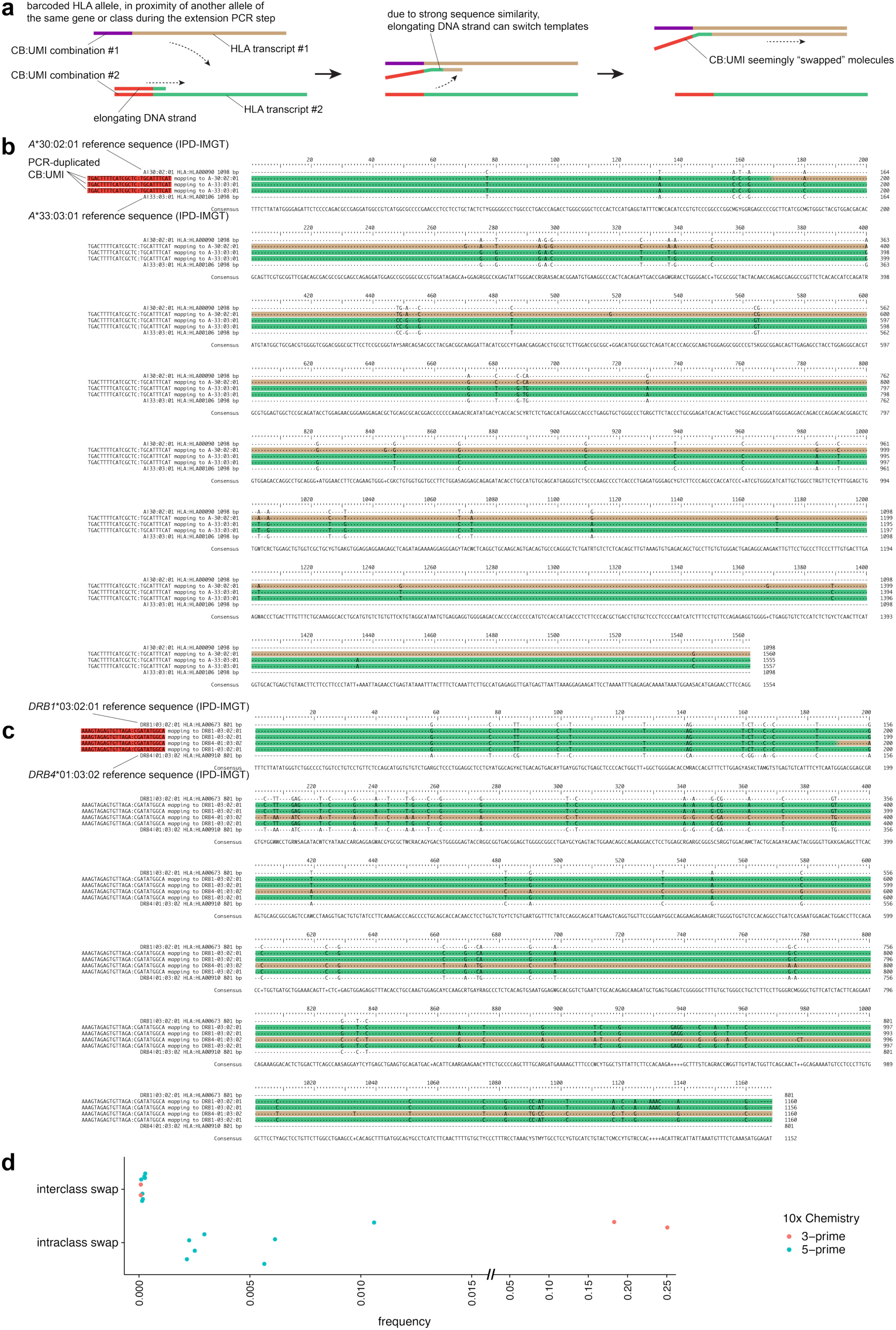
Putative molecular swap mechanism in the case of cell barcodes at the 5’ end of the HLA transcript. **a**, Schematic of the molecular swap process. **b** and **c**, Examples of a molecular swap event impacting 1 of the molecules belonging to the same cell barcode / unique molecular identifier (CB:UMI), highlighting the approximate location at which the elongating DNA sequence switched to a very similar template (to another allele of the same gene in [**b**], and another gene of the same HLA class in [**c**]) represented by the green-to-gold junction. The reference sequences as provided by the IMGT/HLA database are provided to help illustrate. **d**, Molecular swap prevalence depending on whether the swap is between alleles of the same gene or different genes but of the same HLA class (i.e., intraclass swap), or between alleles distantly homologous, e.g., of a different HLA class (i.e., interclass swap).

**Extended Data Fig. 4.**
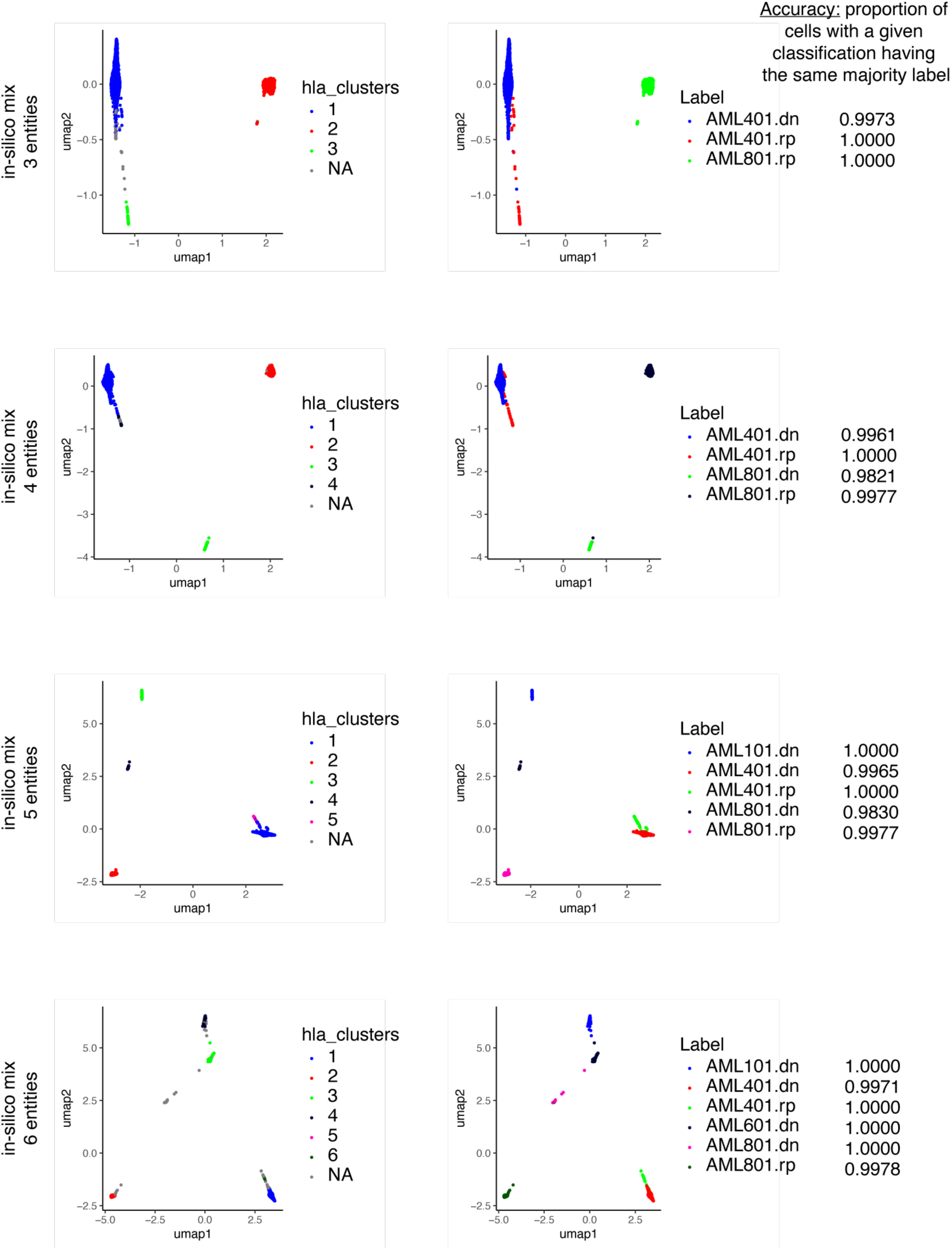
HLA pattern-based classification of cells by scrHLAmatrix from in-silico mixes with 3 or more chimeric entities. The plots show classification of cells based on HLA expression patterns (‘NA’ corresponds to ‘not assigned’) in comparison to their original labeling, known a priori before BAM file merging. The examples shown here are from first guided iteration results. The prediction accuracy, defined as the proportion of cells with a given classification corresponding to the same majority label, is also shown.

**Extended Data Fig. 5.**
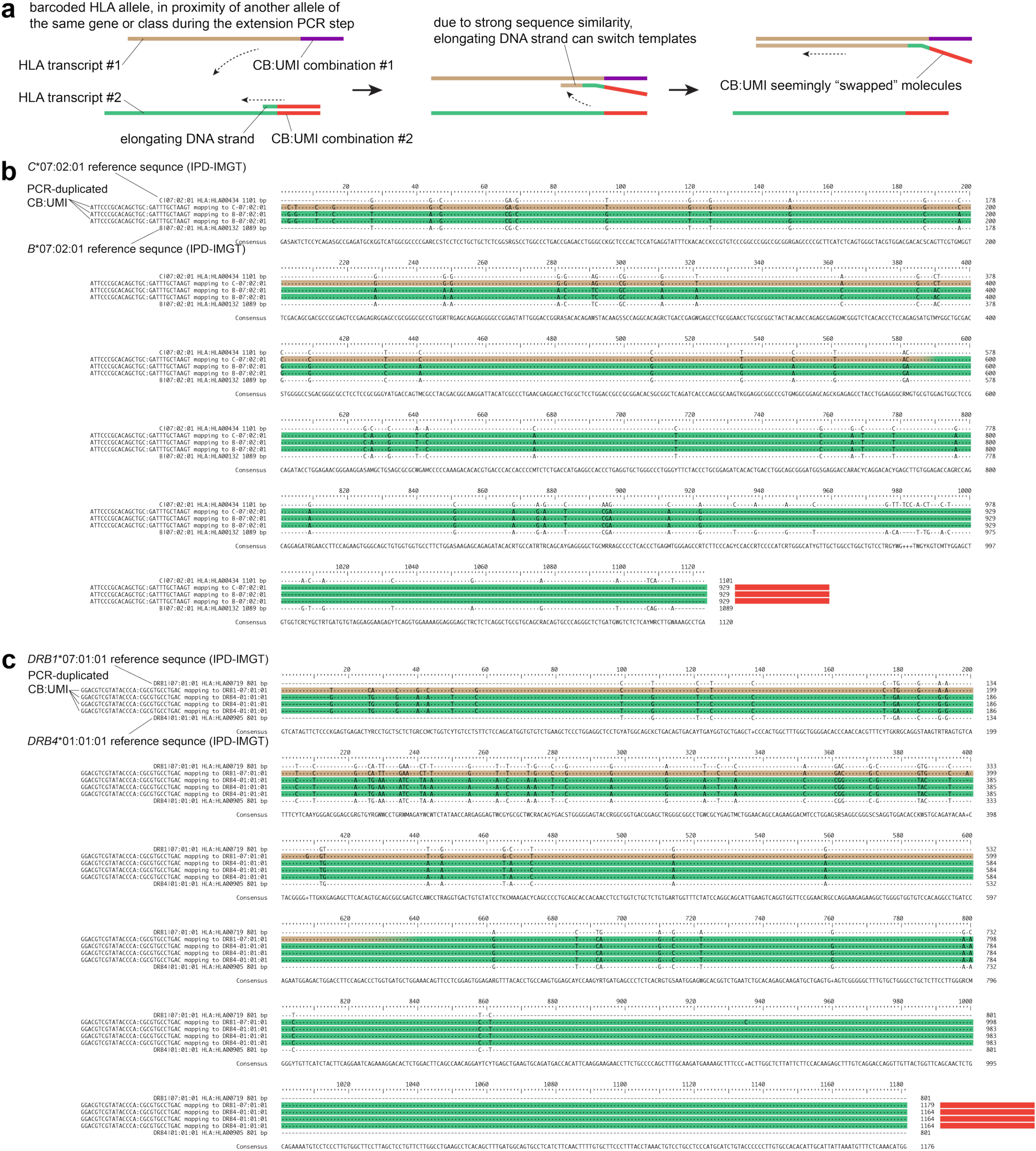
Putative molecular swap mechanism in the case of cell barcodes at the 3’ end of the HLA transcript. **a**, Schematic of the molecular swap process. **b** and **c**, Examples of a molecular swap event impacting 1 of the molecules belonging to the same cell barcode / unique molecular identifier (CB:UMI), highlighting the approximate location at which the elongating DNA sequence switched to a very similar template (to another gene of the same HLA class in [**b**] and [**c**]) represented by the green-to-gold junction. The reference sequences as provided by the IMGT/HLA database are provided to help illustrate.

**Extended Data Fig. 6.**
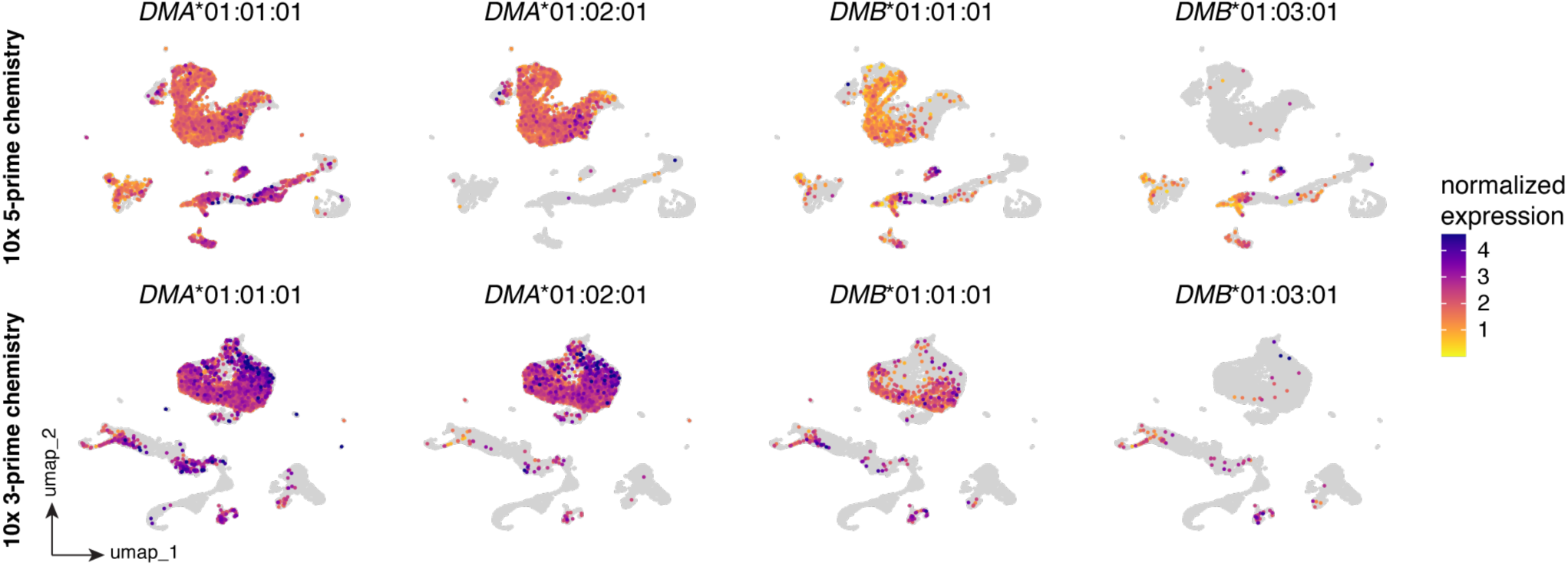
Quantifying the mismatched HLA alleles in samples from the same patient captured in 2 different 10x Genomics chemistries. Normalized expression of *DMA*\*01:01:01, *DMA*\*01:02:01, *DMB*\*01:01:01, and *DMB*\*01:03:01 (*HLA-DMA* and *-DMB* genes were the only genes with allele mismatches between donor and recipient).

**Extended Data Fig. 7.**
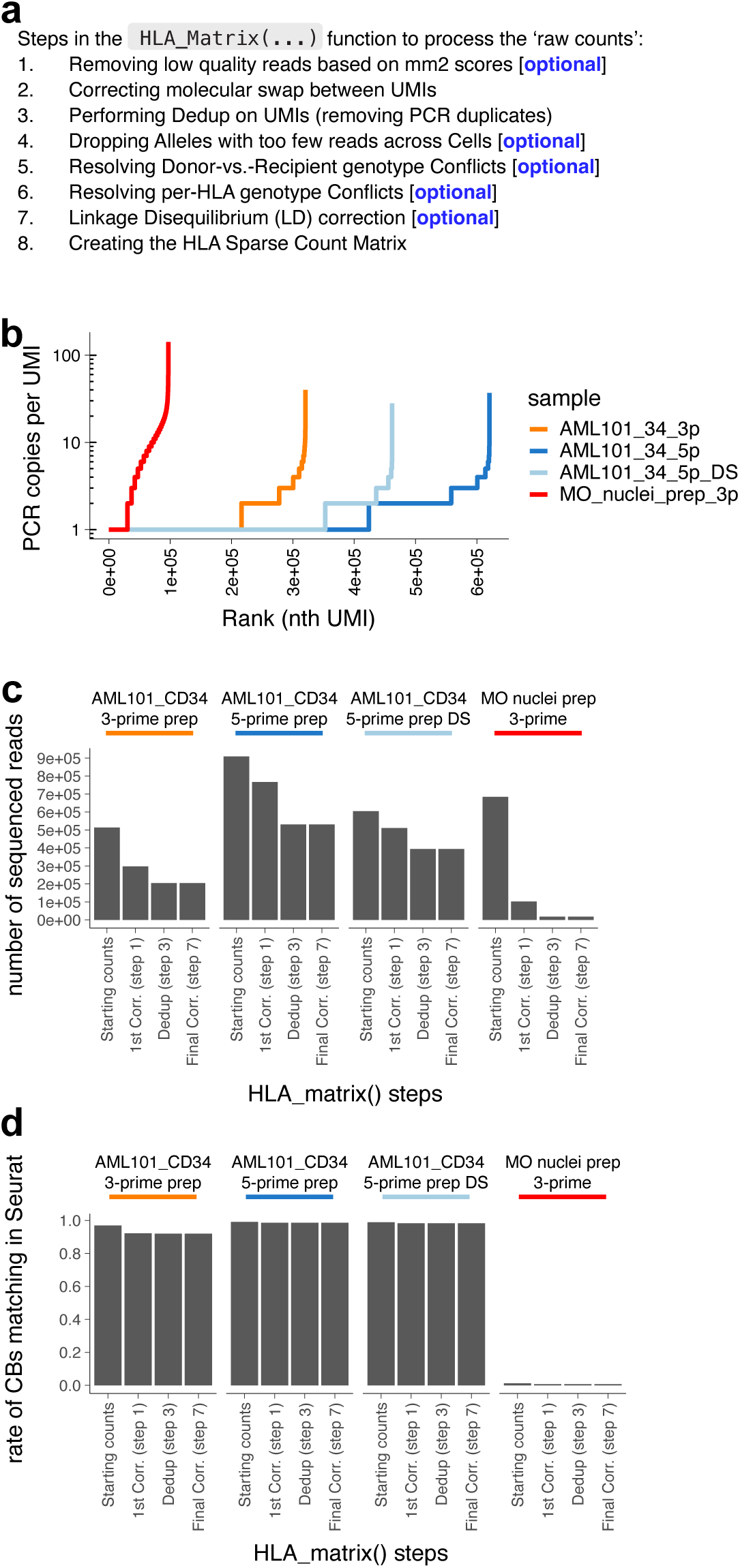
Read count metrics from alignment reads across the various processing steps, comparing 3’ vs. 5’ 10x Genomics capture chemistries. **a**, Processing steps in the making of the sparse numeric matrices from raw read counts. **b**, PCR duplication rates in ‘raw counts’ as an output from scrHLAtag for 4 samples: ‘AML1’ CD34-enriched in 3’ chemistry, ‘AML1’ CD34-enriched in 5’ chemistry, ‘AML1’ CD34-enriched in 5’ chemistry down-sampled (DS) to match the file size from the 3’ chemistry (for a more ‘fair’ at each processing step in the making of sparse numeric matrices from raw read counts. **d**, Rate of cell-dataset-object associated cell barcodes (CBs) matching with the CBs in the read counts at each processing step.

**Extended Data Fig. 8.**
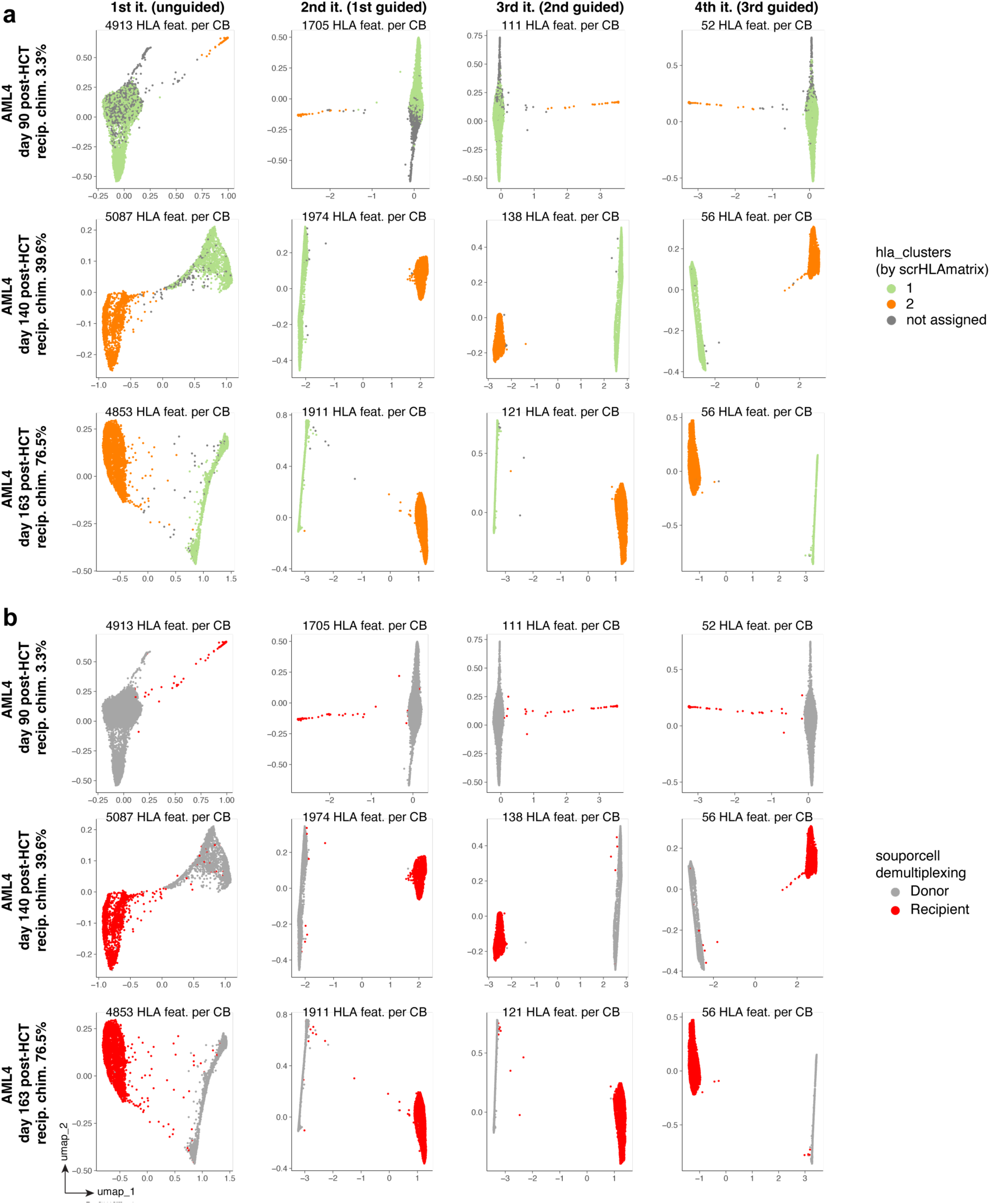
HLA pattern-based classification of cells by scrHLAmatrix. **a**, Classification of cells at 3 timepoints post-transplant, across 4 sequential scrHLAtag iterations. The number of HLA features (i.e., alleles) in the scrHLAtag output is shown for each iteration.

**Extended Data Fig. 9.**
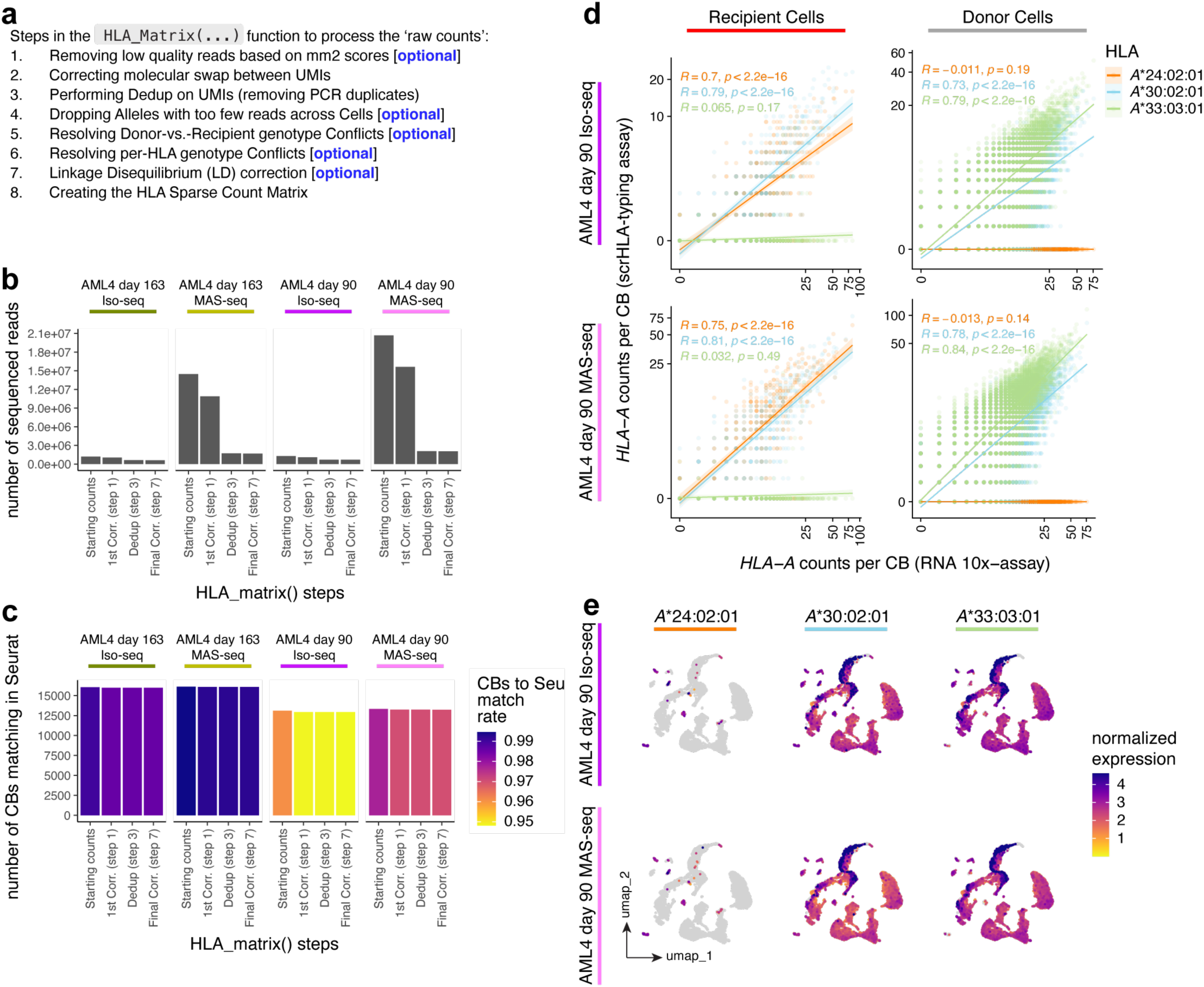
Read count metrics from alignment reads across the various processing steps comparing the standard Iso-Seq method to the augmented MAS-seq method. **a**, Processing steps in the making of the sparse numeric matrices from raw read counts. **b**, Number of reads retained at each processing step. **c**, Number of cell-dataset-object associated cell barcodes (CBs) matching with the CBs in the read counts at each processing step. **d**, correlation of allele counts (UMIs) per CB in the HLA-pulldown assay with the corresponding gene transcript counts per CB in the short-read 10x Genomics RNA assay, in the example of *HLA-A* for ‘AML4’ day 90 post-transplant. **e**, Normalized expression of the 3 alleles of *HLA-A* present in patient ‘AML4’ at the day 90 post-transplant timepoint.

**Extended Data Fig. 10.**
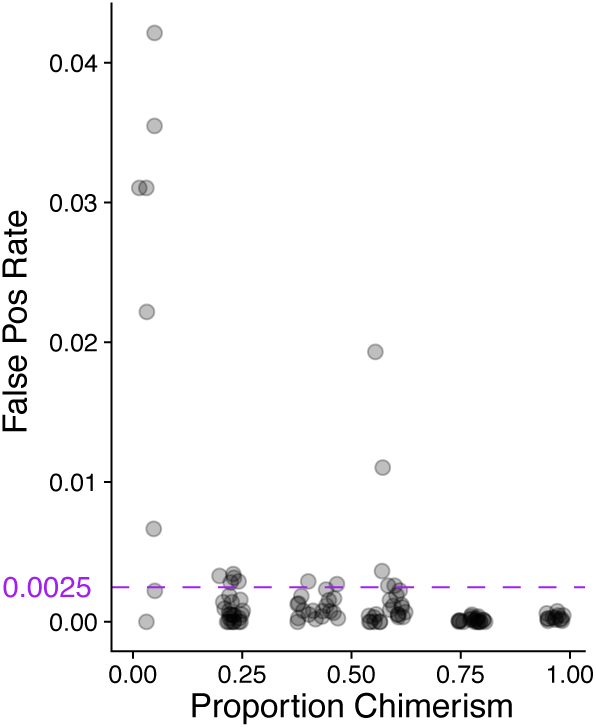
Rates of false positives or out-of-place occurrences (i.e., in reference to souporcell classification, donor-specific alleles in recipient cells and vice-versa, recipient-specific alleles in donor cells) according to the proportion of the analyzed allogeneic entity in the chimeric mix. With a maximum < 5%, the rate rapidly decreases in the chimeric population as its chimerism proportion increases, maintaining a mean rate of 0.25% (purple line).

**Extended Data Fig. 11.**
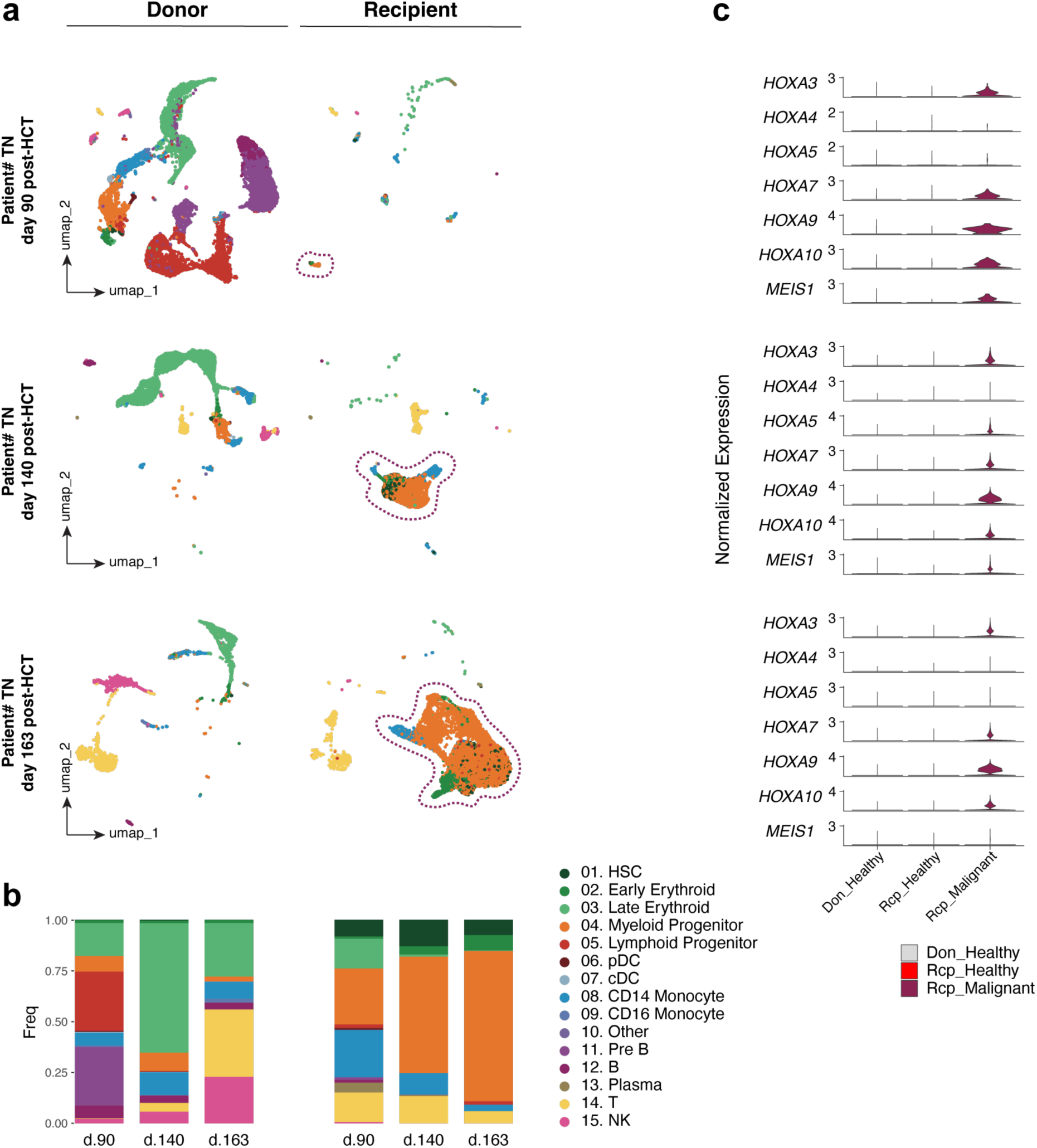
Identifying malignant cells in the AML4 samples across the 3 timepoints post-transplant. **a**, Cell type classification according to viewmastR, split by genetic demultiplexing classification (souporcell). **b**, Cell type composition, split by genetic demultiplexing classification (souporcell). **c**, Identifying candidate cells (clustered together in UMAP space; almost entirely non-overlapping with ‘healthy’ clusters; indicated inside dotted circles in [**a**]) among recipient cells as malignant, and verifying their expression of genes implicated in cellular immortality.

**Extended Data Fig. 12.**
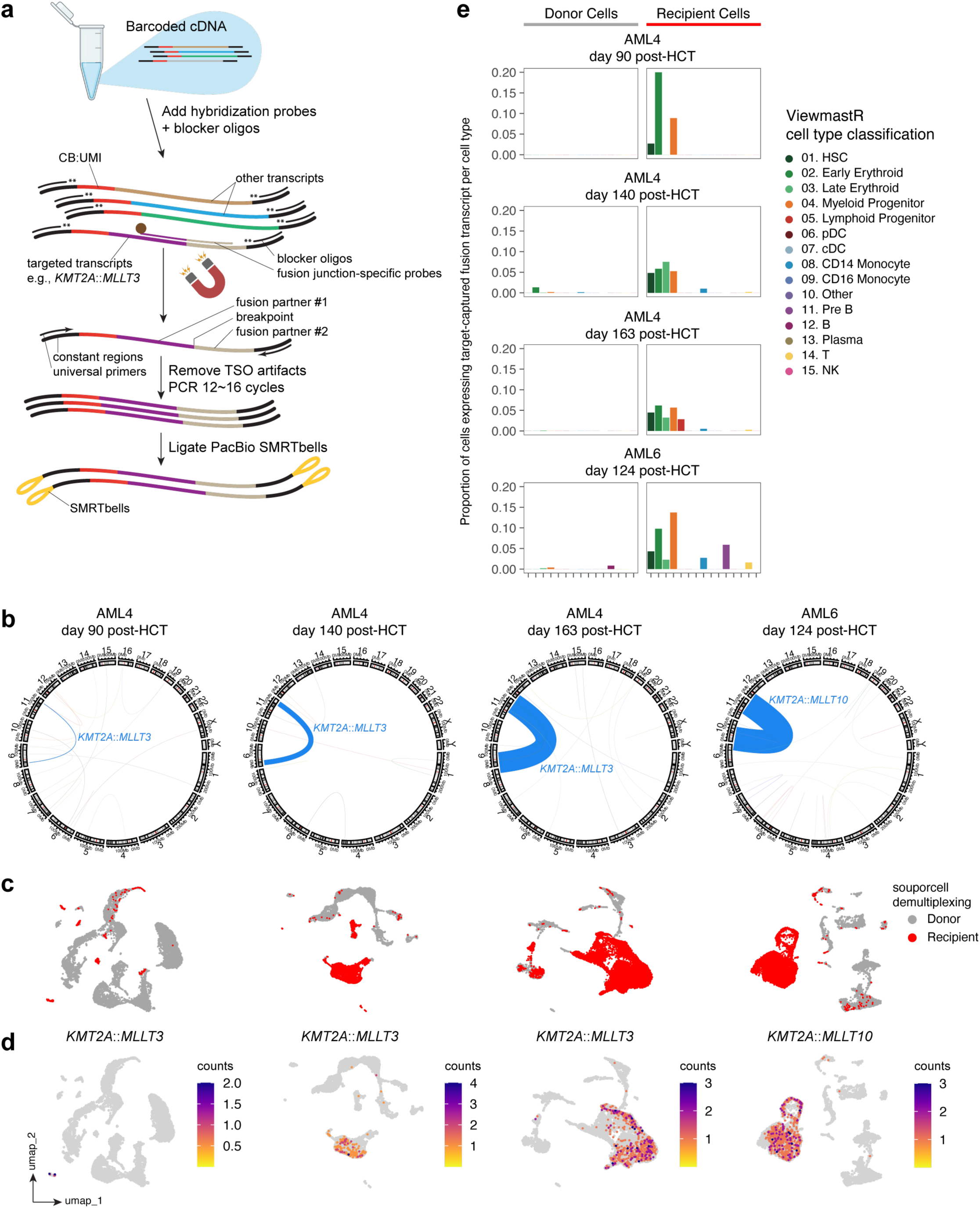
Experimental design and output of targeted gene fusion capture in AML4 and AML6. **a**, Starting product is single-cell barcoded cDNA pools (from a capture platform like 10x Genomics). To that, custom-designed biotinylated hybridization probes (specific to the fusion breakpoint region) and blocking oligos are added. Transcripts with the targeted gene fusions are enriched magnetically (using streptavidin beads), depleted from template switch oligo (TSO) priming artifacts, PCR-amplified, and prepped for long-read sequencing. **b**, Circular plots depicting chromosomal locations (hg38) as rings, and gene fusion partners as colored arcs, starting by the color blue for the fusion event with the highest count (arc thickness proportional to event counts). **c**, Genetic demultiplexing classification (by souporcell) of cells from bone marrow aspirates at days 90, 140, and 163 post-allo-HCT of patient ‘AML4’, and day 124 of patient ‘AML6’. **d**, Expression levels (counts) of the target-captured gene fusion transcripts by single cell. **e**, Proportion of cells expressing target-captured gene fusion transcript per cell type and according to genetic demultiplexing classification (by souporcell); a small proportion of donor cells were positive for fusion expression, dismissed as false positives.

**Extended Data Fig. 13.**
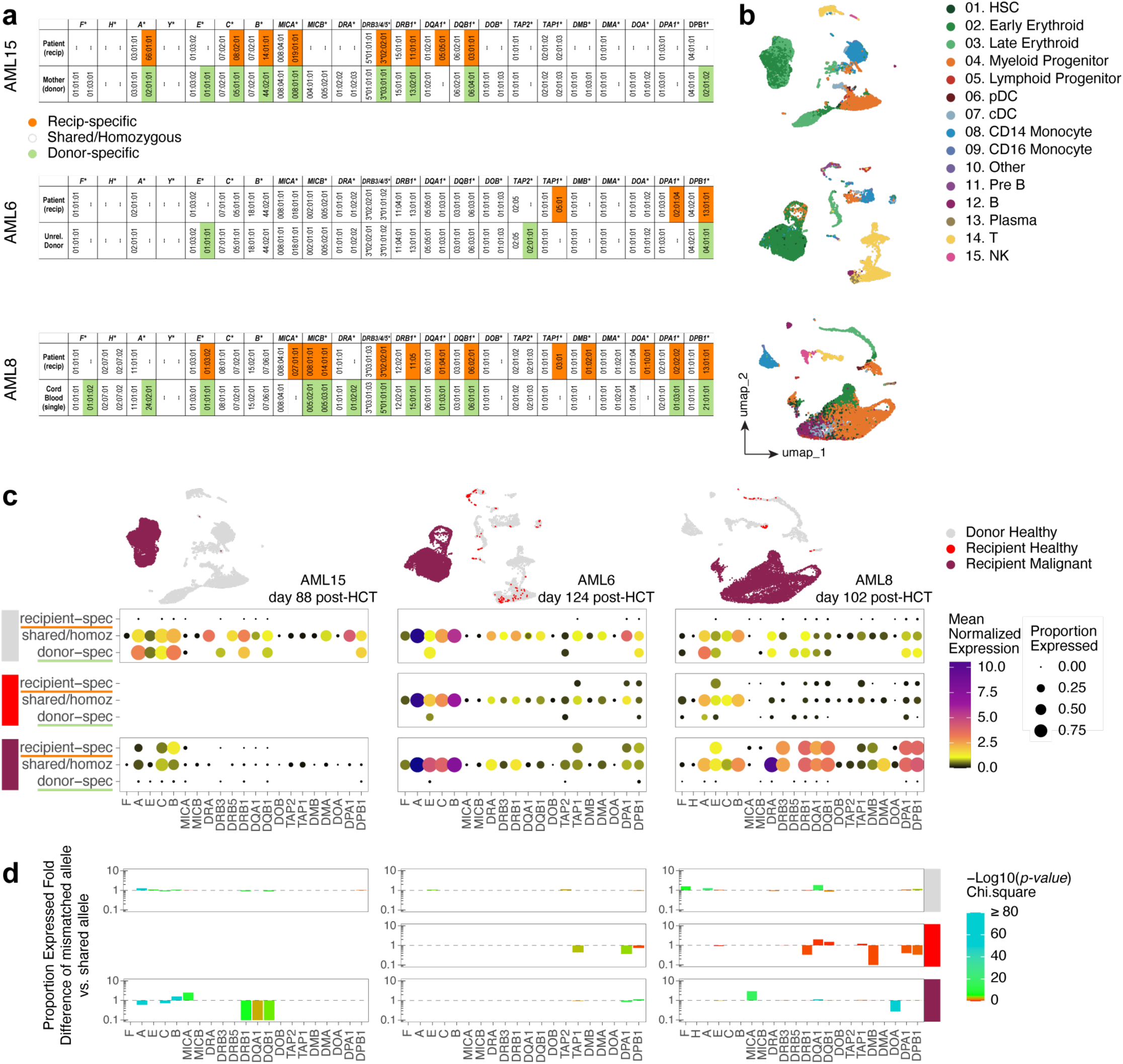
Using additional samples to quantifying allele-specific differential expression across the HLA haplotype in groups of cells. **a**, Genotyping of patients and their donors according to scrHLA-typing. **b**, Cell type classification according to viewmastR. **c**, Mean normalized expression and proportion of cells expressed for the recipient-specific, shared (or homozygous), and donor-specific HLA alleles in the healthy donor, healthy recipient, and malignant recipient cells, for each sample. **d**, Proportion expressed fold change of the mismatched allele (under pressure) vs. shared allele and its statistical significance (Chi-square test), for each sample.

**Extended Data Fig. 14.**
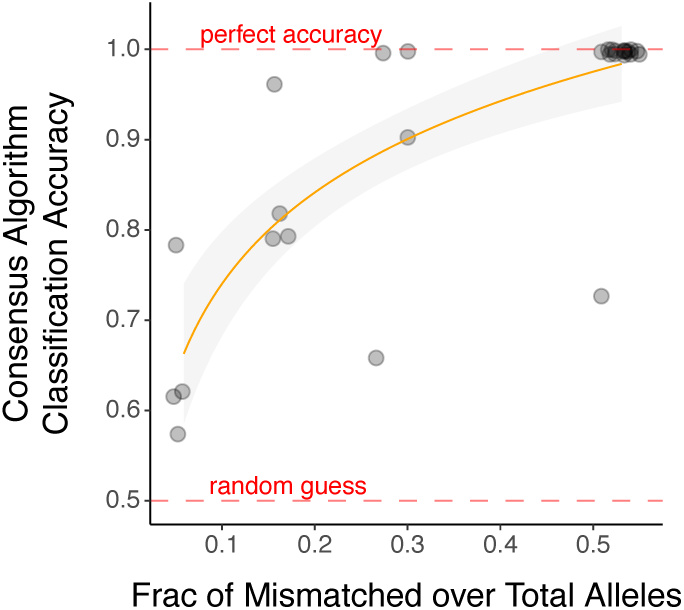
The ability of the scrHLAmatrix classifier, using the consensus algorithm, in correctly predicting the chimeric entities (ground truth: genetic demultiplexing by souporcell) as a function of the abundance of mismatched alleles between the chimeric entities.

**Extended Data Fig. 15.**
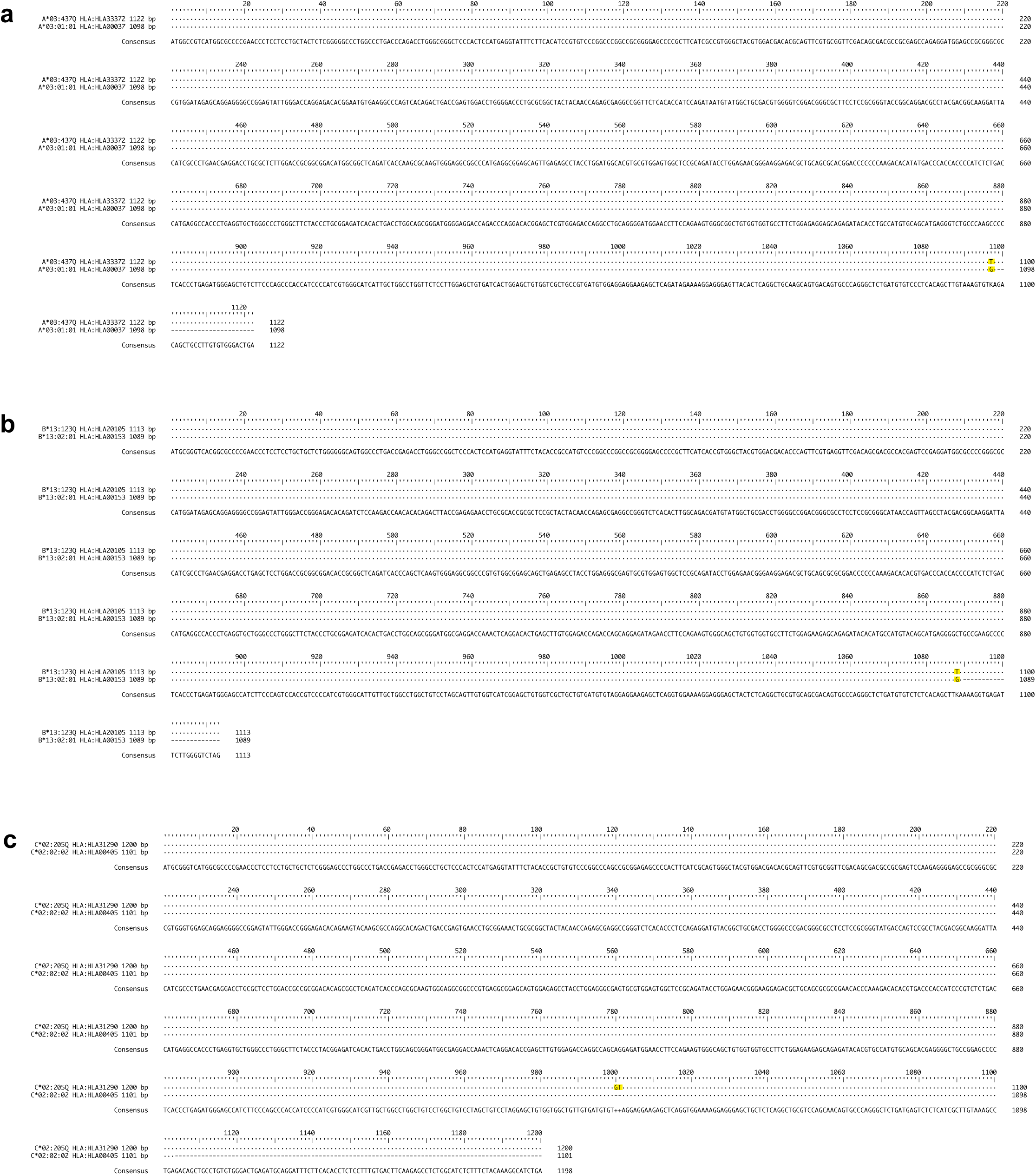
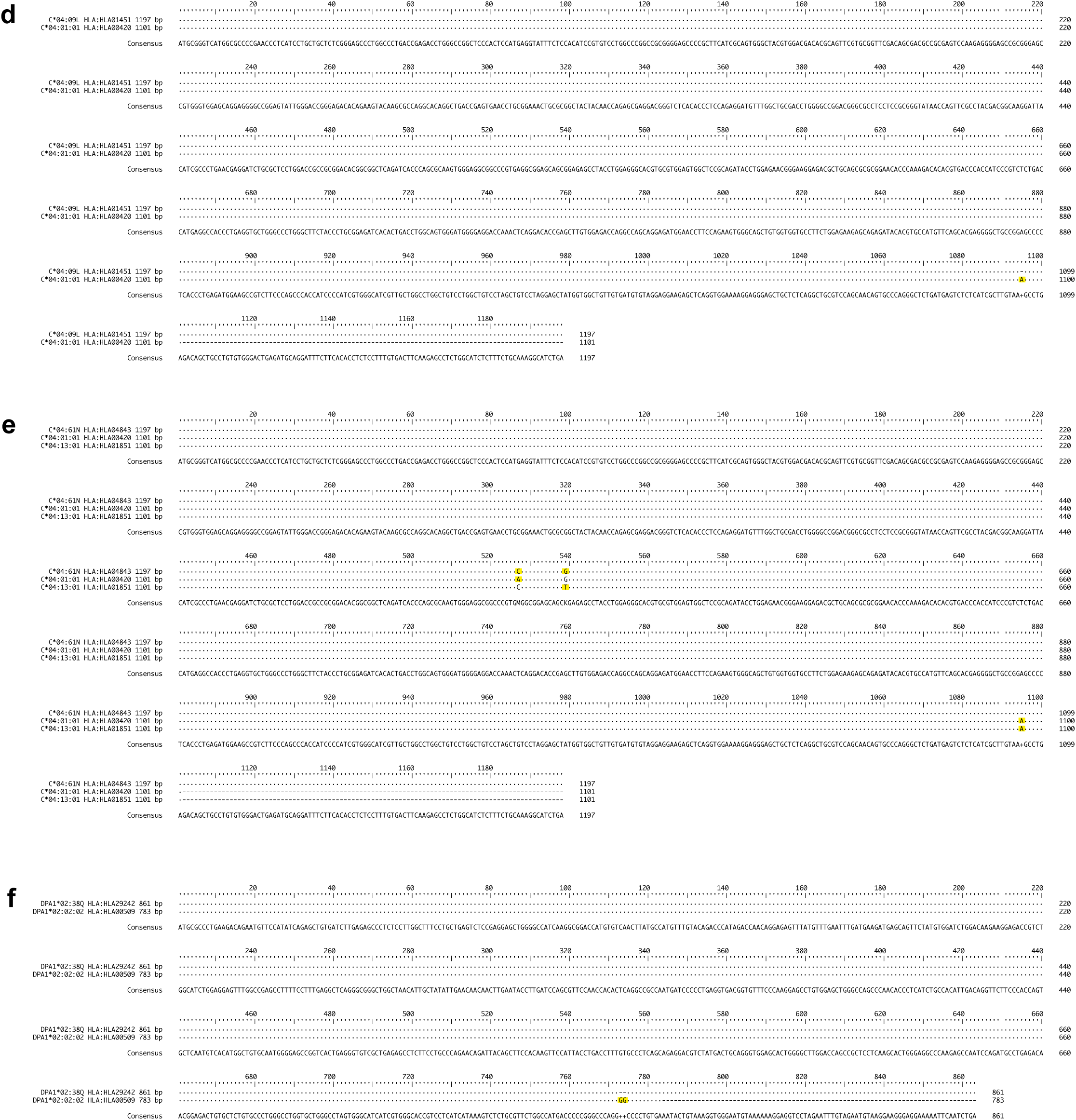
The mRNA/cDNA 3-field reduced IMGT/HLA library contains 6 reference sequences with 24 to 99 additional nucleotides sequenced downstream the 3’ furthest most common sequencing starting position of the remaining references in the library. These references represent alleles: *A*\*03:437Q (**a**), *B*\*13:123Q (**b**), *C*\*02:205Q (**c**), *C*\*04:09L (**d**), *C*\*04:61N (**e**), and *DPA1*\*02:38Q (**f**), to which PacBio long-read sequences belonging to *A*\*03:01:01, *B*\*13:02:01, *C*\*02:02:02, *C*\*04:01:01, again *C*\*04:01:01, and *DPA1*\*02:02:02 will preferentially align, respectively, despite sustaining a 1 to 2 nucleotide(s) mismatch penalty (highlighted in yellow).

